# Parental genomes segregate into different blastomeres during multipolar zygotic divisions leading to mixoploid and chimeric blastocysts

**DOI:** 10.1101/2021.11.05.467317

**Authors:** Tine De Coster, Heleen Masset, Olga Tšuiko, Maaike Catteeuw, Nicolas Dierckxsens, Sophie Debrock, Karen Peeraer, Katrien Smits, Ann Van Soom, Joris Robert Vermeesch

**Affiliations:** Laboratory for Cytogenetics and Genome Research, Department of Human Genetics, KU Leuven, 3000 Leuven, Belgium; Reproductive Biology Unit, Department of Internal Medicine, Reproduction and Population Medicine, Ghent University, 9820 Merelbeke, Belgium; Leuven University Fertility Center, University Hospitals of Leuven, 3000 Leuven, Belgium

**Keywords:** zygote, mitosis, whole-genome segregation errors, chromosomal instability, triploidy, chimerism, mixoploidy, mola, multipolar division, heterogoneic division

## Abstract

The zygotic division enables two haploid genomes to segregate into two biparental diploid blastomeres. This fundamental tenet was challenged by the observation that blastomeres with different genome ploidy or parental genotypes can coexist within individual embryos. We hypothesized that whole parental genomes can segregate into distinct blastomere lineages during the first division through “heterogoneic division”. Here, we map the genomic landscape of 82 blastomeres from 25 embryos that underwent multipolar zygotic division. The coexistence of androgenetic and diploid or polyploid blastomeres with or without anuclear blastomeres, and androgenetic and gynogenetic blastomeres within the same embryo proofs the existence of heterogoneic division. We deduced distinct segregation mechanisms and demonstrate these genome-wide segregation errors to persist to the blastocyst stage in both human and cattle. Genome-wide zygotic segregation errors contribute to the high incidence of embryonic arrest and provide an overarching paradigm for the development of mixoploid and chimeric individuals and moles.

## Background

The haploid sperm and oocyte fuse during fertilization forming a zygote. During the first division of the zygote, both parental genomes segregate into two biparental diploid daughter cells. The genomic constitution of both daughter cells is then maintained through further mitotic divisions (1).

Conflicting with this fundamental tenet, genome-wide (GW) mosaicism is occasionally observed. Co-existence of cells with different genome ploidy or distinct parental genotypes in a single individual are classified as mixoploidy and chimerism, respectively. In a specific form called mosaic GW uniparental disomy, the parental origin of one cell lineage is attributed to one parent only. Chimerism (2–6), mixoploidy (7–15) and mosaic GW uniparental disomy (16–22) have been shown to underlie rare developmental disorders in man and cattle. They are also associated with defined clinical placental manifestations, such as placental mesenchymal dysplasia (23–25), and complete or partial hydatidiform moles in women (26).

Given that mixoploidy and chimerism have been molecularly typed either in patients or placental anomalies, previous analyses rely on the cytogenetic analysis of cells that might have undergone rigorous prenatal selection. Hence, the true mechanistic origin of mixoploidy and chimerism remains elusive. Different models invoking (a combination of) zygotic or polar body fusion, parthenogenetic activation and fertilization or meiotic errors have been put forward to explain their origin (27–36). However, they remain speculative and largely unsupported by molecular or cell biological data.

With the development of concurrent GW single-cell haplotyping and copy number profiling, including haplarithmisis (37,38), mixoploidy and chimerism have been demonstrated at early embryonic stage. It was shown that androgenetic, gynogenetic, biparental and triploid blastomeres can co-exist within individual day-2 and day-3 bovine embryos, both *in vitro* (39,40) and *in vivo* (40). Mosaics were observed in embryos derived from both monospermic, i.e. normal, and dispermic fertilizations. The existence of mixoploidy and chimerism was subsequently confirmed in other bovine (41), and non-human primate (42) *in vitro* fertilized (IVF) cleavage-stage embryos.

Since all cells originate from a single zygote, we hypothesize mixoploidy and chimerism to arise from the segregation of parental genomes into different daughter cells during the zygotic division. We coined this phenomenon “heterogoneic division” and reason that it might be enriched in embryos cleaving into three or four cell directly (multipolar zygotic division) (27,39). In addition, we speculate on different mechanisms that could be a source for heterogoneic division, including the formation of private parental spindles caused by asynchronous parental cell cycles or the formation of a multipolar spindle following the disruption of the gonomeric spindle. That is a common, bipolar spindle in which parental genomes are kept separate on the metaphase plate (43).

To test whether multipolar zygotic divisions coincide with parental genome segregation errors and gain insights in the mechanisms, we analysed the GW haplotype architecture and ploidy state of all blastomeres derived from *in vitro* produced bovine zygotes undergoing multipolar division into three or four blastomeres directly. We created bovine zygotes, recorded their cleavage patterns by time-lapse microscopy and subsequently determined the genomic composition of the blastomeres and blastocysts. Since mixoploidy and chimerism have important clinical implications, we further demonstrate that GW mosaicism also exists in human blastocysts. This analysis proofs that a multipolar zygotic division can result in the segregation of entire parental genomes into separate blastomeres and that resulting GW mosaicism can persist into the blastocyst stage.

## Results

### Genome-wide mosaicism exists in human blastocysts

Considering that bovine and non-human primate embryogenesis parallel human embryogenesis (43– 47), we hypothesized mixoploidy and/or chimerism to be traceable in human cleavage- and blastocyst-stage embryos. For ten embryos, where comprehensive haplotyping-based preimplantation genetic testing (PGT) identified a gynogenetic or androgenetic blastomere at day-3 biopsy, we reasoned that the detected GW aberrations might be indicative of a mixoploid and/or chimeric embryo. Hence, we dissociated the resulting blastocysts for single-cell analysis (Fig. 1A), which revealed a mixture of gynogenetic and biparental cells in two embryos (Fig. 1B). More specifically, for the first embryo, human_E01, comprehensive PGT showed a gynogenetic blastomere. Upon dissociation of the blastocyst, two more gynogenetic cells in conjunction with six biparental cells were retrieved. In the biparental cells, a limited number of (reciprocal) segmental or whole-chromosome losses and gains indicative of mitotic segregation errors were also identified. In addition, one cell with a biparental genomic contribution displayed an abundance of copies of the maternal genome relative to paternal genome and hence, represented a polyploid cell. For the second embryo, human_E02, a clinical gynogenetic biopsy profile with some nullisomies was identified. Analysis of five blastomeres at the blastocyst stage showed the presence of two gynogenetic and three biparental diploid cells. As a result of mitotic segregation errors, segmental or whole-chromosome losses and gains were detected in both the gynogenetic and biparental cells. Taken together, these analyses point to the occurrence of human whole-genome segregation errors and the persistence of these lineages into chimeric and/or mixoploid human blastocysts.

**Fig. 1.**
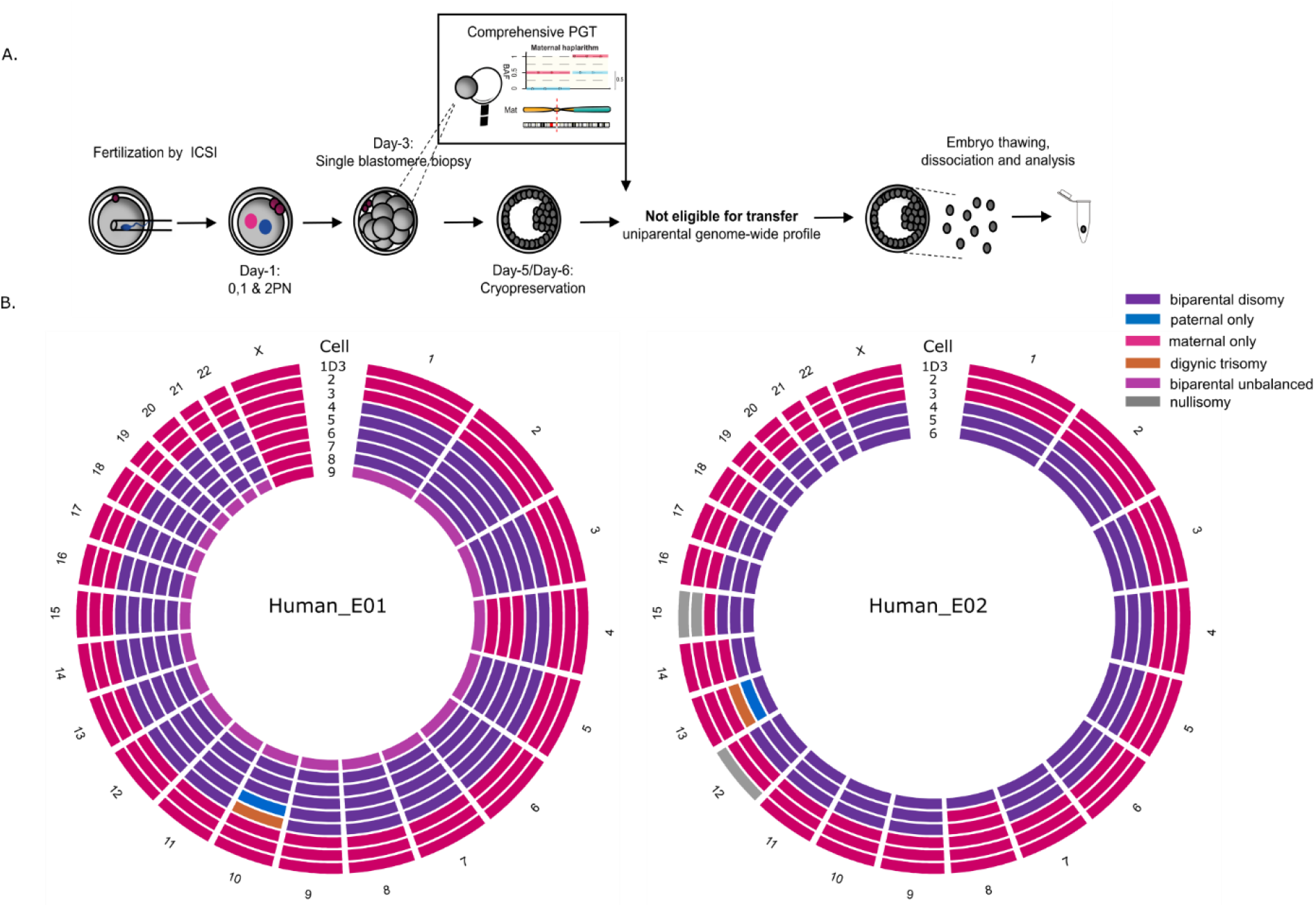
Human blastocysts characterized by GW mosaicism. **A)** Experimental set-up. ICSI: intracytoplasmic sperm injection; PN: pronuclei; PGT: pre-implantation genetic testing. **B)** Circos plots with each circle representing the genome constitution per chromosome of a single cell. The day-3 clinical biopsy (1D3) and cells 2 and 3 are gynogenetic. The remaining cells are biparental diploid or polyploid. Segmental chromosomal errors are not depicted.

### Multipolar zygotic divisions are characterized by whole-genome segregation errors

To pinpoint the origin of whole-genome segregation errors, we analysed the genomic profiles of all blastomeres following a multipolar zygotic division (Fig. 2; S2). A total of 416 *in vitro* produced bovine zygotes were recorded under time-lapse imaging until the zygotic division was observed. Eighty-nine percent of the zygotes cleaved, of which 64% cleaved into two blastomeres and 36% cleaved directly into three or more blastomeres (Movie S1). The average time of the zygotic cleavage was 33.30 hours (ranging from 24.03 to 99.87 hours) after fertilization. Timing of the first division was similar for multipolar and bipolar zygotic divisions (*P* = 0.666).

**Fig. 2.**
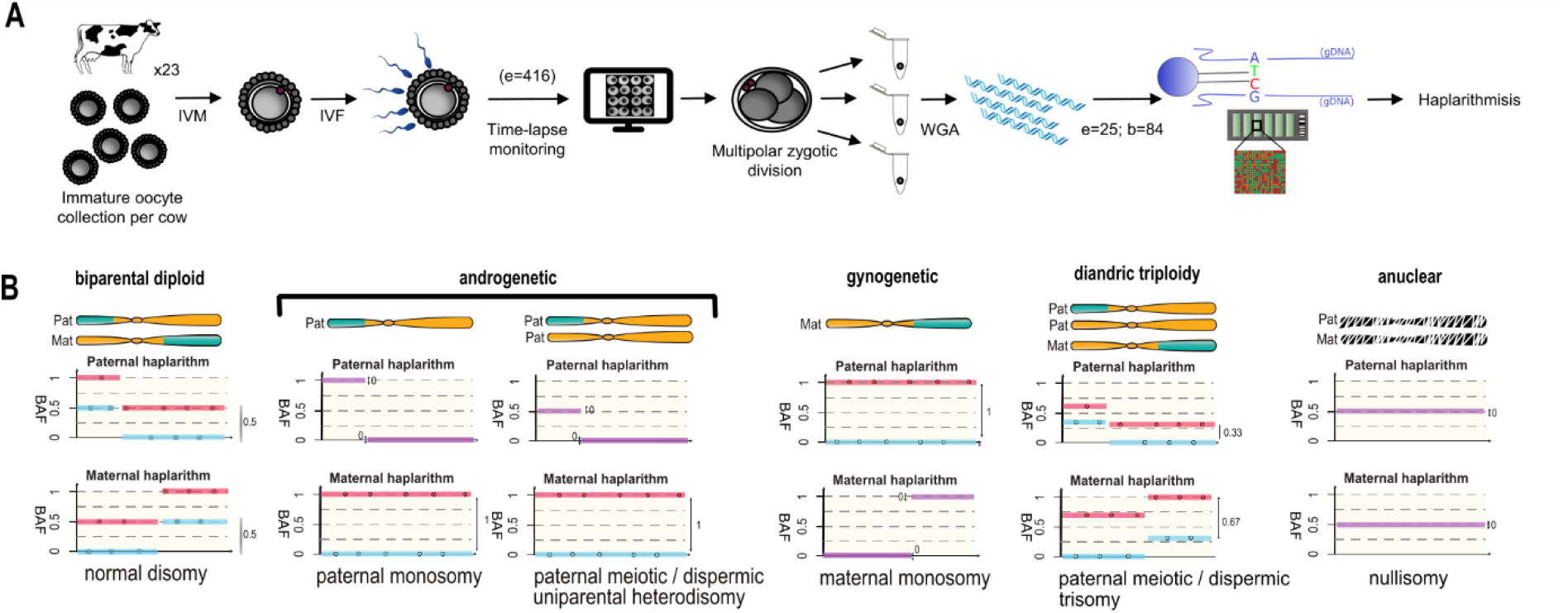
Study design and data analysis by haplarithmisis. **A)** Study design. IVM: *in vitro* maturation; IVF: *in vitro* fertilization; WGA: whole-genome amplification; e=number of embryos; b=number of blastomeres and fragments. **B)** Haplarithm patterns for a selection of genomic constitutions (i.e. normal disomy, paternal monosomy and paternal meiotic/dispermic uniparental heterodisomy) (Full overview in Fig. S2A). Corresponding GW profiles (biparental diploid or androgenetic) are characterized by the manifestation of those patterns throughout the (majority of the) genome. During initial parental phasing, single-cell B allele frequency (BAF) values are assigned to parental informative SNPs, rendering two paternal and two maternal subcategories (blue and red lines). Defined single-cell BAF-values of the segmented subcategories in the sample form haplotype blocks, demarcated by pairwise breakpoints, i.e., homologous recombinations. Haplotype blocks, as well as the distance between the parental SNP subcategories in the paternal and maternal haplarithm, respectively, and the positioning of homologous recombinations, denote the origin and nature of copy number (more detailed explanation in Fig. S2A and (38).

Haplarithmisis was subsequently performed in 82 single blastomeres and two cellular fragments obtained from 25 *in vitro* produced bovine embryos. Embryos cleaved either into three blastomeres (n=16), three blastomeres with a fragment (n=2) or four blastomeres (n=7; Fig. S2). Remarkably, in addition to (segmental) meiotic and mitotic chromosomal aneuploidies, all embryos (n=25) contained GW abnormalities in at least one blastomere, including polyploidy with additional maternal or paternal genomes, uniparental signatures (androgenetic or gynogenetic) or the GW presence of complex aneuploidies. The separate segregation of whole parental genomes in different blastomeres resulted in androgenetic and diploid or polyploid blastomeres, androgenetic and anuclear blastomeres or androgenetic and gynogenetic blastomeres. Such an event occurred in both polyspermic (n=17; 75%) and monospermic embryos (n=3; 12%).

### Anuclear blastomere segregation following multipolar division

In 12 blastomeres and two cellular fragments, the haplarithm plot did not identify any GW haplotype despite a seemingly successful whole-genome amplification (WGA) (Fig. S1; S2). Given that blastomeres contained cell membranes upon visual inspection and had comparable sizes to the other blastomeres, we hypothesized that those cells might not contain nuclear DNA. Hence, we performed low-coverage whole-genome sequencing of the WGA product. For all twelve blastomeres and two cellular fragments, an abundance of mitochondrial DNA was observed (Fig. S1B). In nine blastomeres and two fragments, nuclear DNA was absent. In three blastomeres, a subset of the reads mapped back to some regions in the reference genome, consistent with signals seen on the haplarithm plot, which suggested the presence of chromosomal fragments (Fig. S1C; S2). To exclude external DNA contamination, the mitochondrial sequences were re-assembled and compared, demonstrating each of the sequences to be cow specific (Fig. S1A). Hence, the samples contained mitochondria but no complete nucleus and were categorized as ‘anuclear’. Overall, nine embryos (36%) carried at least one anuclear blastomere and two embryos carried an anuclear cellular fragment.

To confirm and estimate the incidence of anuclear blastomere formation and determine whether anuclear blastomeres can be observed also in bipolar zygotic cleavages, we stained nuclei with Hoechst for a total of 43 and 65 blastomeres, respectively isolated from an additional set of embryos that underwent bipolar zygotic division (n=23) or multipolar zygotic division in three (n=10) or four (n=10) blastomeres. Following multipolar division of the zygote, nine anuclear blastomeres were observed in seven embryos (Fig. S1D). No anuclear blastomeres were observed in bipolar divisions.

### Polyspermic fertilization instigates multipolar division

Based on parental haplotype profiling, 21-embryos (84%) harbored paternal genomes with distinct haplotypes in the zygote prior to multipolar division (Fig. S2). In 18 of these embryos, distinct paternal haplotypes were indicative of polyspermic fertilization (Fig. 3B). In the remaining embryos (E08, E24, E25), one or multiple identical GW paternal homologous recombination sites in combination with the GW heterodisomy made it impossible to distinguish fertilization by multiple sperm from fertilization by a single diploid sperm, caused by a paternal meiotic error (Fig. 2B; S2; (48,49)).

**Fig. 3.**
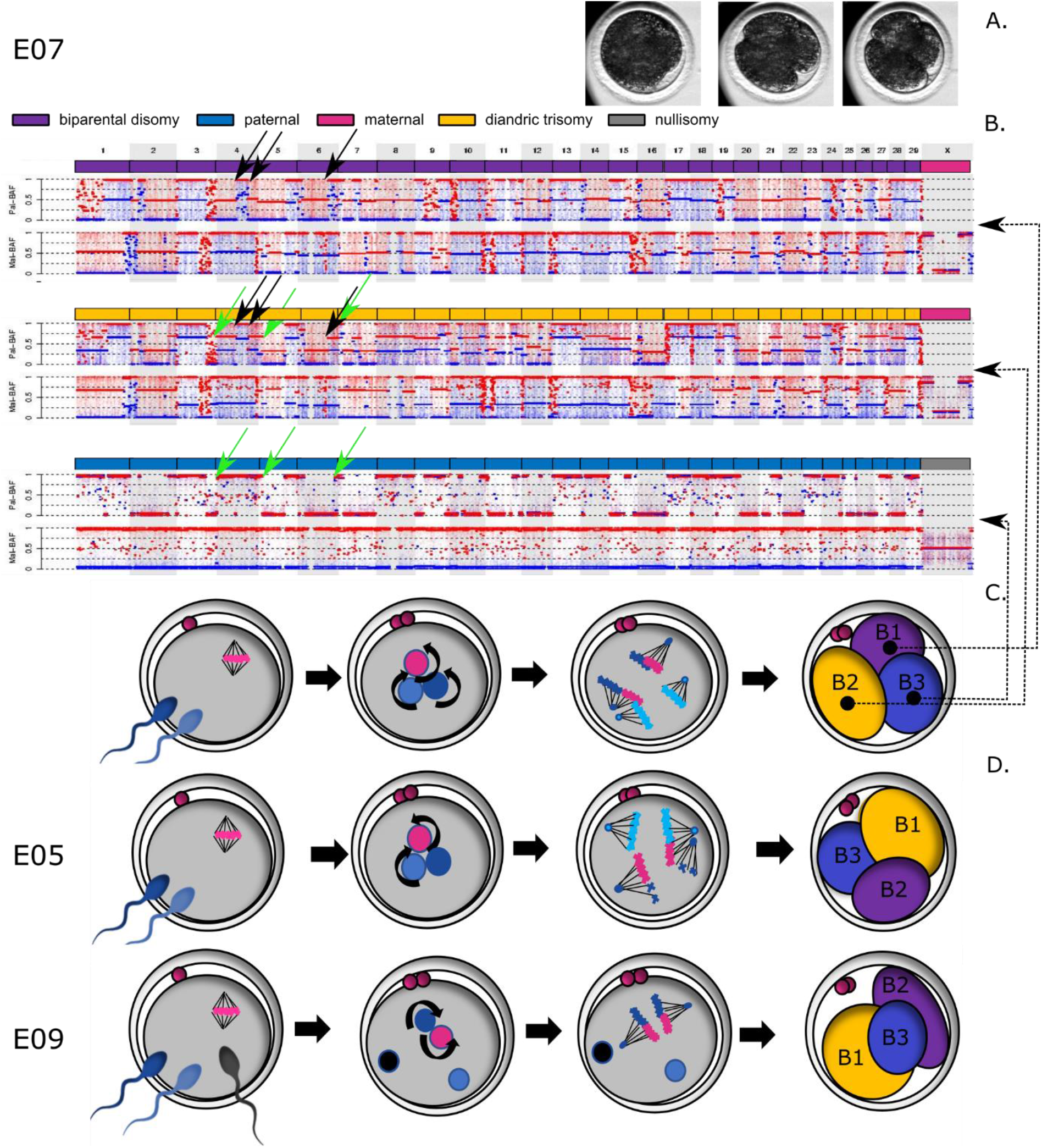
Haplarithmisis unveils GW mosaicism following multipolar zygotic division. Curved arrows depict replication of the genome. **A)** Three chronological time-lapse images of E07 (initiation of the cleavage furrow, the ongoing first division and the embryo immediately after cleavage and before cell isolation of the cleaving zygote). **B)** Haplarithm profiles for the biparental diploid (B1), the diandric triploid (B2) and the androgenetic blastomere (B3). For each blastomere from top to bottom, respectively, we depict the GW interpretation per chromosome and the paternal and maternal haplarithms (See Fig.2 and S2A for detailed explanation). The GW paternal haplarithms uncovered different homologous recombination sites in the biparental diploid (black arrows) and the androgenetic blastomere (green arrows) and a combination thereof in the diandric triploid cell pointing towards dispermic fertilization. **C)** Schematic representation of the likely mechanistic origin, including replication and pronuclear apposition of the parental genomes and karyokinesis by a tripolar spindle leading to a multipolar division of the zygote. **D)** Schematic representation of possible events leading to the segregation of a zygote in biparental diploid, a diandric triploid and an androgenetic cell lineage in two other embryos.

In contrast to abundance of paternal genome, an additional maternal genome was identified only in one embryo (E04). Based on haplotype profiling, presence of the extra maternal genome was due to a meiotic error. Whole-genome maternal meiotic errors in three other embryos were also characterized by the loss of large segments (E16) or the complete loss of the maternal genome (E01 and E10) (Fig.S2). This suggests that polyspermic fertilization is the main driver of multipolar division.

### Segregation errors occur via different mechanisms

We classified embryos in five categories, based on blastomere haplotype profiles (Fig. 2B; S2). The first three categories were consistent with the segregation of a whole parental genome to a distinct cell lineage during the zygotic division, i.e. heterogoneic division. Mapping the segregational origin of the genomic content revealed distinct mechanisms to cause parental genome segregation during multipolar zygotic division.

#### 1. Embryos with diandric triploid, biparental diploid and androgenetic blastomeres

Three embryos (12%) consisted of an androgenetic, a biparental diploid and a diandric triploid blastomere (Fig. 3; S2). Despite the similar ploidy constitution, the molecular mechanisms preceding segregation were different. Specifically, in two of these embryos (E05 and E07), two distinct paternal haplotypes were present, providing evidence that the oocyte was fertilized by two sperm. Both paternal haplotypes were present in the triploid blastomere and one of the other blastomeres (Fig. 3B; S2). Hence, a tripolar spindle likely tethered the maternal and both paternal genomes during the first mitotic division (Fig. 3C,D). In embryo E07, two copies of all three genomes were present. Therefore, replication of the maternal and both paternal genomes likely preceded the formation of a mitotic tripolar spindle (Fig. 3B, C). In contrast, embryo E05 contained only one copy of one of the paternal genomes. Hence, one paternal pronucleus was not replicated before the onset of mitosis, but was stochastically incorporated into the hypodiploid and hypotriploid sister blastomeres, respectively, resulting in reciprocal paternal losses and gains for several chromosomes (Fig. 3D; S2).

In E09, three distinct paternal haplotypes were present, indicating that the embryo was fertilized by three sperm. One of the paternal haplotypes was shared between the diandric triploid and the biparental diploid blastomere. A second and a third paternal haplotype were identified in the diandric triploid and androgenetic blastomere, respectively. Hence, one of the paternal pronuclei and the maternal pronucleus replicated and participated in the karyokinesis, while two other paternal genomes were ejected by a cytoplasmic protrusion into the diandric triploid and androgenetic blastomeres (Fig. 3D; S2).

#### 2. Embryos with biparental and androgenetic blastomeres

Thirteen embryos (52%) consisted of a combination of biparental and androgenetic blastomeres (Fig. 4A,B; S2). Multiple paternal haplotypes were detected in all embryos, indicating polyspermic conceptions. In three embryos, fertilization occurred by more than two sperm.

**Fig. 4.**
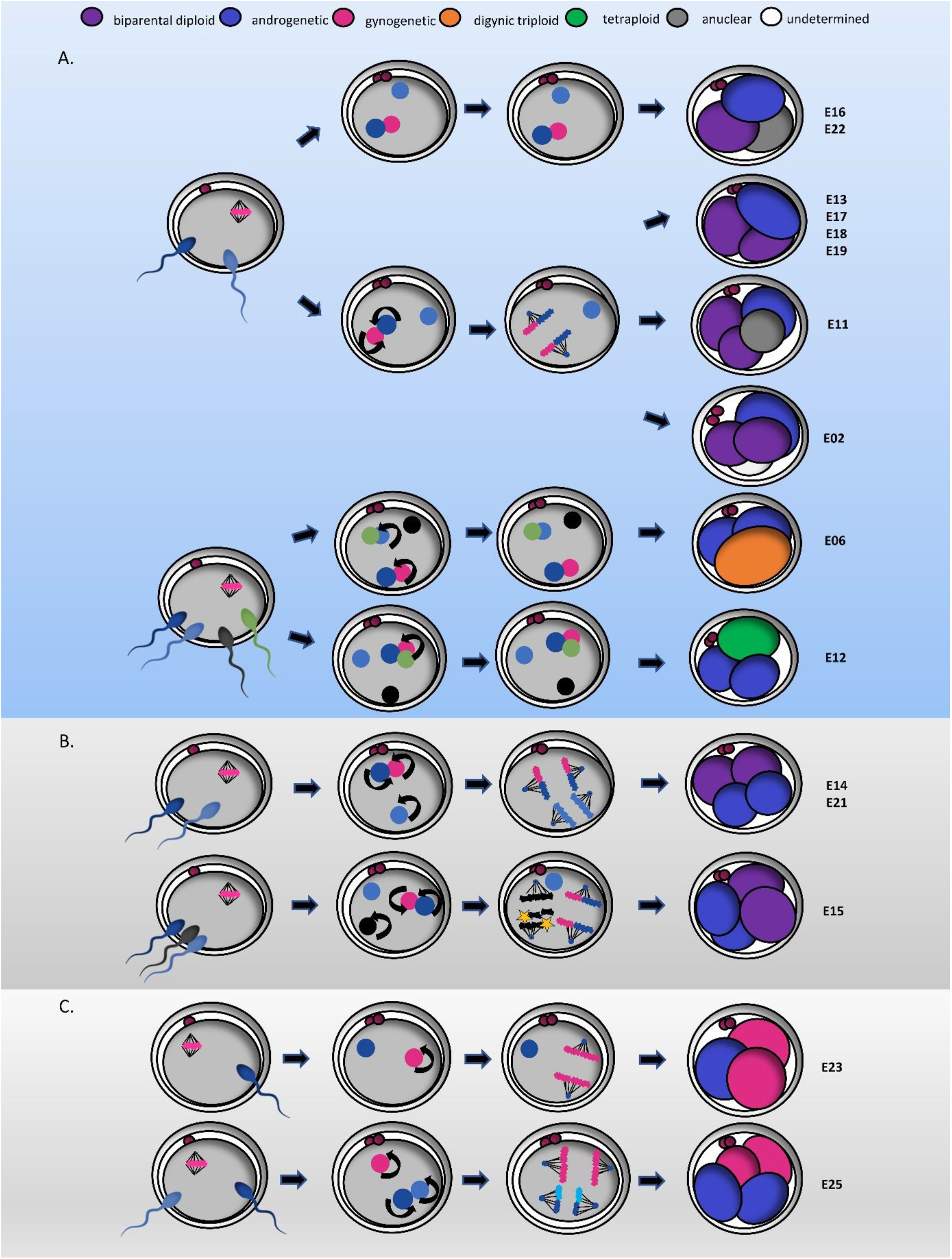
Likely events leading to the segregation of a zygote into biparental and androgenetic (A, B) or androgenetic and gynogenetic blastomeres (C). Curved arrows depict replication of the genome. In **A)** cytoplasmic protrusion of paternal genome(s) is depicted following polyspermic fertilization. In some embryos, cytoplasmic protrusion occurred in parallel with the replication and karyokinesis of the primary maternal and paternal genome. In E06 and E12 some of the parental genomes were replicated but failed to undergo karyokinesis. In **B)** segregation of an additional paternal genome occurred by a second, paternally organized spindle. In E15, a third paternal genome was protruded to one of the androgenetic blastomeres and genome-wide chromosomal losses occurred on one paternal genome (yellow stars). In **C)** private parental spindles are established around the maternal and the paternal genomes (E25) following polyspermic fertilization, or only around the maternal genome following normal fertilization (E23). In E23, the paternal genome is expelled in a separate blastomere by cytoplasmic protrusion.

In embryos with two biparental diploid blastomeres, the presence of the same maternal and paternal haplotypes and occasional reciprocal aneuploidies in both biparental blastomeres of some embryos (chromosome 8 in E13, chromosome 9 in E19 and complex in E21) confirmed that both parental genomes were segregated via one spindle (Fig. 4A,B; S2).

The androgenetic blastomere(s) in all 13 embryos presented one or more paternal haplotypes that were distinct from the haplotype observed in the biparental blastomere(s) (Fig. S2). Hence, the extra paternal genomes might have segregated into a separate blastomere either by cytoplasmic protrusion (Fig. 4A), or by the operation of a second spindle that organized the second paternal pronucleus in an ectopic metaphase plate (Fig. 4B). The latter was recognized by the presence of two androgenetic blastomeres containing the same paternal haplotype, indicating that a second spindle segregated the chromatids (Fig. S2). The presence of a segmental deletion of chromosome 3 in one and duplication in the other androgenetic blastomeres of E21 confirms that the segregation occurred following genome replication (Fig. S2).

In some embryos, additional complexities were found (Fig. 4; S2). In one androgenetic blastomere of E06, regions of heterodisomy throughout the genome indicated the presence of three paternal copies of two distinct haplotypes. This might have occurred via cytoplasmic protrusion of one replicated and one non-replicated paternal pronucleus to the same androgenetic blastomere. The simultaneous presence of a digynic triploid blastomere with two identical maternal copies in the same embryo, pointed towards the apposition of two parental nuclei, but replication of only the maternal nucleus before the zygotic division. In E12, four paternal haplotypes were present of which two were included in the tetraploid biparental cell with balanced parental genomes containing only one maternal haplotype. Hence, fertilization occurred by four sperm and two paternal genomes were ejected by cytoplasmic protrusion. The other paternal genomes and the replicated maternal genome remained in the tetraploid blastomere. Regions of heterodisomy throughout the genome in an androgenetic blastomere of E15 hinted to the involvement of an inactive, third paternal nucleus during the segregation of the second paternal pronucleus by the ectopic spindle. Moreover, lost chromosomes throughout the genome of the other androgenetic blastomere suggested the partial loss of the paternal nucleus segregated by the ectopic spindle. In E16, (segmental) maternal chromosomes were missing throughout the genome, likely due to a meiotic error.

#### 3. Embryos with androgenetic and gynogenetic blastomeres

In two zygotes (8%), the parental genomes segregated into three or four blastomeres, each containing a maternal or paternal genome (Fig. 4C; S2). In E25, regions of heterodisomy in both androgenetic blastomeres indicated fertilization by two haploid or a single diploid sperm followed by genome replication and karyokinesis by two spindles. Each spindle segregated the paternal or maternal genome into two androgenetic or gynogenetic blastomeres (Fig. 4C; S2). Embryo 23 contained only a single androgenetic and two gynogenetic blastomeres. Hence, the paternal genome was likely segregated by cytoplasmic protrusion rather than by a second spindle.

#### 4. Androgenetic embryos

In two embryos (8%), the blastomeres did not contain a maternal genome. The zygotes cleaved in three or four blastomeres consisting of two anuclear and one or two androgenetic blastomeres, respectively (Fig. S2). Both androgenetic blastomeres in E10 contained an identical haplotype. Hence, they resulted from genomic replication of the paternal genome followed by karyokinesis. In E01, a single androgenetic and two anuclear blastomeres were observed which pointed towards the cytoplasmic protrusion of the paternal pronucleus.

#### 5. Polyploid embryos

Two embryos (8%) consisted of two polyploid and one anuclear blastomere (Fig. S2). The polyploid blastomeres were diandric triploid (E24) and tetra-andric pentaploid or triandric tetraploid (E03). Regions of heterodisomy indicated polyspermy in both embryos. Based on genomic profiles that were identical (E24) or semi-identical containing multiple reciprocal aneuploidies (E03), the segregation of all genomes likely occurred by a single bipolar spindle. In E03, reciprocal aneuploidy resulted in diandric trisomies in one blastomere and tetra-andric hexasomies in the other. As the diandric trisomic regions contained two identical paternal haplotypes, polyspermic fertilization followed by two replications of one of the paternal genomes is thought to have caused the presence of two copies of the same paternal genome and one copy of a different paternal genome. Additional cross-over sites in the tetra-andric pentasomic regions pointed towards the presence of an additional paternal genome in the third blastomere. A combination of replication and segregation errors in two of the three paternal genomes might have caused the complex embryonic profile.

One embryo (E08) consisted of a triandric tetraploid blastomere and two anuclear blastomeres (Fig. S2). Regions of heterodisomy disclosed the presence of three distinct paternal haplotypes in the tetraploid blastomere. Therefore fertilization occurred by three sperm, followed by cytokinesis and either a non-synchronized karyokinesis or no karyokinesis at all.

##### Other profiles

Embryo E04 (Fig. S2), with two nuclear and one anuclear blastomere, was fertilized by one sperm. In the nuclear blastomeres, a digynic triploid profile was likely a result of an erroneous first meiotic division due to the non-extrusion or reabsorption of the first polar body in the zygote. Visualisation of two polar bodies in the time-lapse video, indicated that non-extrusion of the maternal genome into the polar body was more likely. In addition, complex, non-reciprocal maternal and paternal chromosomal losses were observed throughout the genome of one blastomere resulting in diploid biparental chromosomes, maternal heterodisomy and monosomies of maternal or paternal origin. Those errors may have taken place during the first mitotic division.

Embryo 20 presented a complex mosaic chromosome profile (Fig. S2). Three blastomeres contained the GW presence of multiple, seemingly random chromosome aneuploidies. Alternating (segmental) paternal disomies and maternal monosomies or disomies were found in a first blastomere, and alternating biparental chromosomes and maternal monosomies were found throughout the genome of a second blastomere. A third blastomere showed an androgenetic profile with segmental losses throughout the genome. Different recombination patterns in the regions where a paternal haplotype was simultaneously present in different blastomeres pointed towards dispermic fertilization. The profile would be consistent with the replication of three parental genomes followed by a tripolar spindle segregating the chromosomes randomly in three blastomeres.

### Genome-wide abnormalities resulting from heterogoneic division persist in the blastocyst-stage bovine embryo

To prove that GW aberrant blastomeres can propagate and persist following heterogoneic division, we cultured an additional cohort of bovine embryos to the blastocyst stage following multipolar zygotic division and dissociated and genotyped eight to 12 single cells of six blastocysts (Fig. 5A; Fig. S3). The subgroup of sampled cells presented a biparental diploid constitution in two blastocysts (E26, E31) and a GW mosaic constitution in four blastocysts (E27 to E30). In addition, a number of meiotic (E26) and (reciprocal), (segmental) mitotic aneuploidies (E28, E30, E31) were retrieved. In the four GW mosaic embryos, cells of the same blastocyst showed distinct paternal haplotypes or GW paternal heterodisomy, confirming polyspermy to instigate multipolar zygotic division (Fig. 5C). Specifically, three of the GW mosaic blastocysts contained biparental diploid cells in the embryonic mass and one or two androgenetic blastomeres of a distinct paternal haplotype in the perivitelline space (E27, E29, E30) (Fig. 5B, C). One blastomere of E29 presented with complex chromosomal losses, resulting in a fragmented genome containing (segmental) androgenetic and gynogenetic chromosomes and nullisomies. The fourth chimeric blastocyst (E28) contained diandric androgenetic cells with GW regions of heterodisomy in the embryonic mass and one biparental diploid cell with complex (segmental) paternal losses in the periphery (Fig. 5B, C).

**Fig. 5.**
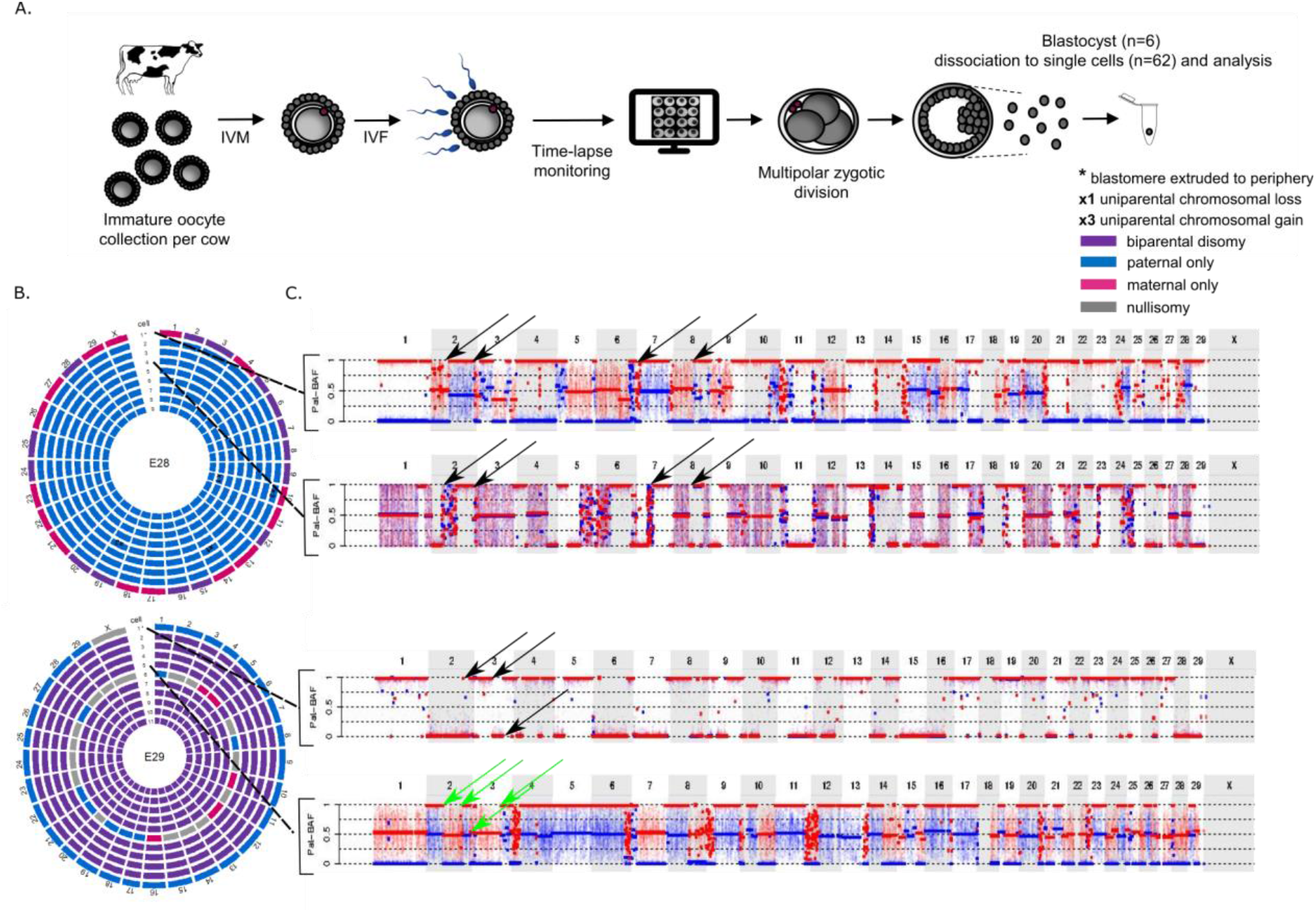
Genome-wide mosaicism persists to the bovine blastocyst stage. **A)** Experimental set-up. IVM: *in vitro* maturation; IVF: *in vitro* fertilization. (Full overview in Fig. S3). **B)** Circos plots of two bovine blastocysts in which each circle represents the genome constitution per chromosome of a single cell. In E28, cell 1 presented with alternating biparental disomic and maternal chromosomes while the remaining cells were androgenetic. In E29, cell 1 was found to be androgenetic while cells 2 - 5 and 7 - 11 were biparental diploid. Cell 6 displayed alternating paternal chromosomes, maternal chromosomes or nullisomies. **C)** Paternal haplarithms of two cells from each embryo. In E28, identical homologous recombination sites were retrieved in the androgenetic and diploid cells (black arrows). Additional heterodisomic regions throughout the genome of cells 2-9 (Pat-BAF = 0.5) revealed the presence of a second paternal haplotype. In E29, the androgenetic and diploid cells present with different paternal homologous recombination sites (green arrows). Segmental chromosomal errors are not depicted.

## Discussion

### Heterogoneic division occurs during the first zygotic division

The co-existence of diploid, triploid and uniparental cell lineages was discovered in bovine and non-human primate pre-implantation embryos (39–42). Here, we demonstrate this occurs in human embryos as well. It was hypothesized that GW mosaic embryos are caused by heterogoneic division, during which the parental genomes in the zygote are segregated into distinct cell lineages (27,39). By tracing division kinetics and determining the genomic structure of all single blastomeres, we provide direct evidence that heterogoneic division is common in embryos that undergo multipolar zygotic division.

### Heterogoneic divisions occur via a multitude of pathways

By reconstructing the segregational origin of the parental haplotypes in the daughter blastomeres, we obtain evidence that heterogoneic division is caused by distinct mechanisms, which mainly perturb the formation of one bipolar spindle. We provide indirect evidence for tripolar, ectopic paternal and private parental spindles and pronuclear extrusion of paternal genomes. Some of these mechanisms have been proposed in theoretical models (29,39,43).

Parental genomes are located at spatially distinct places on the metaphase plate in different species (50–55). Segregation of parental genomes to distinct blastomeres has recently gained a biological foundation, as spindles in murine and bovine zygotes were shown to form around each parental genome (53,54). In a normal zygotic division, these so-called “dual spindles” align and fuse during early metaphase, resulting in a single bipolar spindle before genome segregation. However, faulty alignment of one of the spindle poles would result in the independent segregation of a whole parental genome. Indeed, experimental induction of spindle non-alignment gave rise to segregation of parental genomes in different directions leading to gross mitotic aberrations (e.g. formation of binucleated blastomeres or multipolar division) (53). In addition, experimental non-unification of parental genomes led to formation of private parental spindles, each segregating only one of the parental genomes (55).

All zygotes undergoing multipolar zygotic division reported here contained whole-genome segregation errors and the majority were polyspermic. This confirms previous observations that polyspermy induces multipolar zygotic divisions (56–60). In humans and other non-rodent mammalian zygotes, the sperm centrioles nucleate microtubuli which, upon fertilization, mediate pronuclear movement for the apposition of the male and female genome (55,61,62), DNA clustering at the pronuclear interphase (55) and spindle formation (63,64). During fertilization with more than one sperm, additional pronuclei and centrosomes may form additional spindles. According to the dual spindle model (53,54), more than two spindles around the parental pronuclei might align and fuse resulting in balanced karyokinesis of all parental genomes. Such segregation has been suggested to occur in a fraction of tripronuclear (3PN) human embryos by cytogenetic studies (29) and was evidenced by two bovine embryos in our cohort, consisting of two polyploid blastomeres (Fig. S2; E03, E24). In contrast, non-alignment of one of the additional spindles at one pole could give rise to a tripolar spindle allocating distinct maternal and paternal genomes to distinct blastomeres at the first cleavage. Two mixoploid embryos consisting of androgenetic, biparental diploid and diandric triploid blastomeres are consistent with this model (Fig. 3, S2; E05, E07). Tripolar spindles have been observed before in human dispermic zygotes (65,66). In case of non-alignment at two spindle poles, a tetrapolar spindle would be formed. This tetrapolar spindle may comprise of an “ectopic paternal spindle” that segregates the additional paternal genome into androgenetic blastomeres, alongside the primary spindle segregating the biparental diploid blastomeres. (Fig. 4B; S2; E14, E15 and E21). Alternatively, the tetrapolar spindle comprises of two “private parental spindles” segregating the maternal and two paternal genomes into gynogenetic and androgenetic blastomeres, which is consistent with one embryo in our cohort (Fig. 4C; S2; E25). The physiological occurrence of ectopic paternal and private parental spindles is evidenced by visualization of ectopic metaphases and spindles in 3PN bovine, human and rhesus monkey zygotes (60,63,67–71). Also in presumed normally fertilized zygotes, maternal meiotic and a sperm-derived mitotic-like spindles have been observed to coexist in the same oocyte by Van Blerkom and colleages (72) in so-called ‘silent fertilizations’, i.e. fertilizations in which no pronuclei are visualized in the zygote. More recently, live-staining experiments confirmed the frequent coexistence of private maternal and paternal spindles in the same zygote in normally fertilized bovine zygotes (54). We observed one normally fertilized embryo which was consistent with the formation of a private maternal spindle (Fig. 4C; E23).

In most embryos, however, we demonstrate that excessive paternal genomes were not segregated by a spindle but were expelled to one or more distinct cell lineage(s), resulting in an androgenetic or polyploid blastomere and one or more biparental diploid blastomeres (Fig. 3D, E09; 4A; S2). This observation confirms early cytogenetic studies on human 3PN embryos which predicted the exclusion of one haploid genome from the metaphase plate of the first division, resulting in 2n, 2n/3n mosaics and 1n/2n derivatives (29,56,66,68,69,73–76). Based on cytogenetic analysis after the first cleavage (56,68) of human 3PN zygotes such an exclusion was suggested to occur by a “pronuclear extrusion” of a haploid genome, i.e. a pseudo-cleavage in which cytokinesis occurs while the genome is still packed inside the sperm head or in the pronuclear stage. More recently, such pronuclear extrusion of a parental genome was visualized in polyspermic first cleavage bovine embryos (70).

### Other whole-genome segregation errors

Some GW aberrations were caused by processes other than heterogoneic division. Among those, three embryos were likely the consequence of a meiotic error resulting in the gain of an distinct maternal copy (Fig. S2; E04) or the complete loss of the maternal genome (Fig. S2; E01, E10). For the latter two embryos, the complete androgenetic genome was consistent with the segregation of the maternal genome to a polar body. Formation of androgenetic zygotes by the segregation of the maternal genome to the polar body can be induced in mice by mutagenesis of meiotic genes such as protein 1 (*Mei1*), responsible for double-stranded break formation (77).

Another embryo with one polyploid nuclear blastomere points to a dissociated karyo- and cytokinesis (Fig. S2; E08). It has been suggested that multiple parental genomes, spindles and centrosomes, all together forming a complex spindle apparatus, may impede timely karyokinesis (78,79). Alternatively, cytokinesis may also occur without karyokinesis, resulting in multinucleation of the polyploid blastomere and two anuclear blastomeres. Such multinucleated blastomeres have been reported in other human (79–82) and bovine (83) studies.

### Anuclear blastomeres

Unexpectedly, cytokinesis often occurs without concurrent genome segregation resulting in anuclear blastomeres. These have been shown to exist in human, primate, porcine and bovine early cleavage-stage embryos using microscopy and sequencing (42,79,82–85). How these anuclear blastomeres originate remains unknown, but as confirmed by our single-cell haplotyping, sequencing and nuclear staining results, they have been associated with polyspermy and multipolar division (83,84). Some suggested that dissociation of the karyokinesis from the cytokinesis, due to altered spindle dynamics or altered actin polymerization/depolymerization related due to *in vitro* maturation could underlie anuclear blastomere formation (79,84). However, direct evidence is lacking at present. Alternatively, we suggest that centrosomes may be involved in the segregation of anuclear blastomeres as they can prematurely detach from the sperm head (62,86), dissociate and migrate away from the spindle in the cytoplasm resulting more often in abnormal cleavage kinetics (54) and are able to initiate cytokinesis independently (87).

### Mixoploidy and chimerism occur in human

Different lines of evidence suggest that heterogoneic division occurs also in human embryos. The first, indirect evidence was provided by early cytogenetic studies that reported mixoploid embryos following presumed polyspermic fertilization (29,56,66,69,73–76) or normal fertilization (88–90) in human cleavage-stage and blastocyst-stage embryos. Current GW genotyping methodologies identified large scale aneuploidy during early development (91,92), but most are unable to detect the ploidy nor parental origin of aneuploidy in a single assay. With the recent adoption of single-cell GW haplotyping methods in PGT, the existence of androgenetic, gynogenetic and triploid profiles in human blastomere and trophectoderm biopsies has been uncovered (49,93–96). Moreover, mixoploidy and/or chimerism was recently inferred from single-cell transcriptome data in human blastocysts (97). Here, we directly show GW aberrations, identified in biopsied cleavage-stage embryos, to persist throughout the preimplantation period forming mixoploid and/ or chimeric human blastocysts.

A second line of evidence comes from the observation that multipolar zygotic divisions occur frequently (8.3-26%) in human embryos (98). In human, multipolar divisions of embryos have been associated with tripolar spindles generated by centrosome dysfunction or dysregulation caused by variants of the Polo-like kinase 4 (PLK4) gen(99,100). These result in complex chromosomal losses rather than the segregation of whole parental genomes trough heterogoneic cell division. While GW segregation errors observed here were mostly instigated by polyspermy, the development of polyspermic embryos following IVF in human reproductive medicine is avoided by the selection against multi-pronuclear zygotes. Yet, heterogoneic division is not necessarily restricted to polyspermic fertilization as evidenced by E23, genome-wide mosaic human blastocysts and earlier results (39). Also, some of the proposed mechanisms, such as private parental spindles, have been described in normally fertilized embryos (54,72). Yet, it remains unknown whether zygote formation in general and more specifically, dual spindle formation, proposed as the basis for heterogoneic division, is conserved between bovine and human (101,102).

Outgrowth or survival of one or more of the blastomeres following heterogoneic division likely contributes to low human fecundity and provides an overarching explanation for the persistence of GW anomalies in fetuses and patients (Fig. 6) (27). Multipolar zygotic divisions have been linked to impaired embryonic development, implantation and pregnancy rates in both human(98,103–107) and bovine (58) embryos. Also our time-lapse data reveal a reduced blastocyst rate from bovine zygotes that present with a multipolar division (20.9 ± 0.04%; n = 78) compared to zygotes that present with a normal division (35.4 ± 0.03%, n = 277) (*P* = 0.02, unpublished data). Following heterogoneic division, androgenetic, gynogenetic, biparental and triploid blastomeres could have a different evolution: some might propagate better than others and perhaps, undergo a selective advantage. As both maternal and paternal transcripts make an important contribution to preimplantation development (108), it can be assumed that polyploid, haploid and uniparental diploid blastomeres, resulting from whole-genome segregation errors, have a selective disadvantage compared to normal diploid blastomeres. Similarly to the complex aneuploid embryos, GW mosaic embryo constitution may therefore cause developmental arrest by an insufficient number or lack of diploid cells and the occurrence of gene dosage imbalances following genome activation (49,109) (Fig. 6A). On the other hand, compensatory proliferation of the diploid blastomere and/or active selection against the polyploid, haploid or uniparental diploid could result in the progressive depletion of blastomeres with whole-genome errors and ensure a sufficient number of viable diploid blastomeres (109,110) (Fig. 6A). By analysing several cells from bovine blastocysts, in specific after multipolar zygotic division, and from human blastocysts, we demonstrate that the resulting bovine and human blastocysts can contain different genomic constitutions. It seems that biparental diploid cells indeed have a developmental benefit and that cells with GW abnormalities, such as an androgenetic and complex aneuploid cells are selected against by extrusion to the perivitelline space. Also others have reported on the exclusion of complex aneuploid or uniparental blastomeres during blastocyst development (42,96,98). Nonetheless, some embryos did retain the GW mosaic or androgenetic embryo constitution within the blastocyst proper. We hypothesize that, occasionally, polyploid, haploid and uniparental diploid cells are viable and can develop into rare pre- and post-natal and placental aberrations dependent on the selective pressures at play. These include chimeric and mixoploid cell populations, occasionally observed in patients with developmental disorders, sesquizygotic twinning (111) and molas (Fig. 6B,C).

**Fig. 6.**
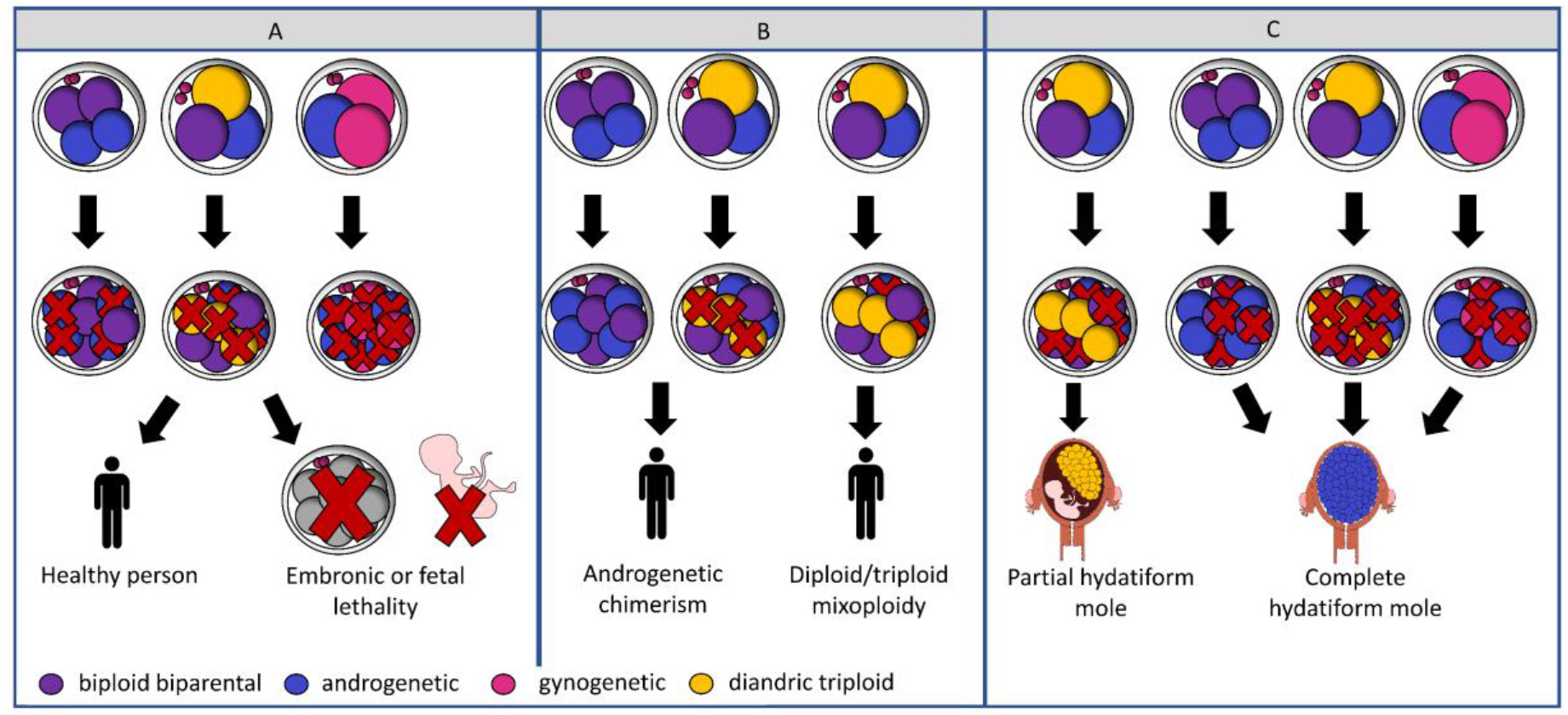
Model explaining the origin of mixoploidy, rare chimeric individuals and clinical placental outgrowths. Following heterogoneic division, the different blastomeres of the mosaic embryos may develop differently.

## Conclusions

In conclusion, this study generated direct proof of heterogoneic division in mono- and polyspermic bovine zygotes and showed that different underlying mechanisms result in a variety of chimeric and mixoploid embryos. In addition, GW segregation errors were found to persist to the human and bovine blastocyst. The proposed mechanisms of heterogoneic division are indirectly supported by recent microscopic observations. The models challenge current cell biological dogmas, including the mechanisms triggering cell division, zygotic spindle formation and the existence of nuclei at different stages of the (nuclear) cell cycle. By combining novel live-imaging technologies with single-cell genome and transcriptome analysis of embryos it will become possible to unravel the cascade of cellular and molecular events involved in the meiotic to mitotic conversion of the zygote and deviations from the normal process that result in whole-genome segregation errors.

## Methods

### Bovine embryo *in vitro* production (IVP)

#### Media and reagents

Basic Eagle’s Medium amino acids, Minimal Essential Medium non-essential amino acids, TCM-199-medium, Ca^+2^/Mg^+2^-free PBS, kanamycin and gentamycin were purchased from Life Technologies Europe (Merelbeke, Belgium) and all other components were obtained from Sigma-Aldrich (Schnelldorf, Germany), unless otherwise stated. All the media were filter-sterilized using a 0.22 μm filter (GE Healthcare, Chicago, US) before use.

#### Procedure

Bovine embryos were produced by routine *in vitro* methods (112). Briefly, bovine ovaries from Belgian Blue cows (*Bos taurus*) were collected at the local slaughterhouse and processed within 2 h. The ovaries were washed three times in warm physiological saline solution supplemented with kanamycin (25 mg/mL). Follicles between 2- and 8-mm diameter were punctured with an 18G needle and a 10mL syringe in 2.5 mL of Hepes-Tyrode’s albumin-pyruvate-lactate (Hepes-TALP) and kept separate per ovary. Cumulus-oocyte-complexes were collected using a stereomicroscope, washed in Hepes-TALP and washed in maturation medium, which consisted of modified bicarbonate-buffered TCM-199 supplemented with 50 μg/mL gentamycin and 20 ng/mL epidermal growth factor. Subsequently, maturation occurred per donor in 500 μL maturation medium in four-well plates (Nunc™) for 22 h at 38.5°C in 5% CO_2_ in humidified air. For fertilization, frozen-thawed semen of three single Holstein-Friesian bulls (*Bos taurus*) at a concentration of 1 × 10^6^ spermatozoa/mL was used following separation of spermatozoa over a discontinuous Percoll gradient (45% and 90%; (GE Healthcare, Chicago, US)). Fertilization was achieved by incubating the matured oocytes with spermatozoa for 21h at 38.5°C in 5% CO_2_ in humidified air. Specifically, E01-E20 were fertilized with semen of bull A, E21-E25 were fertilized with semen of bull B and E26-E30 were fertilized with semen of bull C. The presumptive zygotes were transferred to synthetic oviductal fluid (SOF) supplemented with essential and non-essential amino acids (SOFaa), 0.4% BSA, and ITS (5 μg/mL insulin, 5 μg/mL transferrin, and 5 ng/mL selenium) after removal of excess spermatozoa and cumulus cells by a 3 min vortex step. One or two Primo Vision™ micro well group culture dishes (Vitrolife, Göteborg, Sweden) per donor cow were used. These dishes are designed by the well-of-the-well (WOW) principle, consisting of 9 or 16 small wells covered by a 40 μL droplet of medium and 3.5 mL mineral oil to prevent evaporation. In each WOW dish, 9 or 16 presumed zygotes of the same cow were placed in individual wells under a time-lapse imaging system and incubated at 38.5°C for eight days in a trigas incubator (5% CO2, 5% O2 and 90% N2).

#### Time-lapse imaging system

The WOW dishes were placed into the sample holder of a compact, digital inverted microscope (Primo VisionTM; Vitrolife, Göteborg, Sweden) which was placed in the incubator. The focus was set mechanically and all embryos were positioned in the microscopic field-of-view. Every 5-10 min, a single picture was taken. Using time-lapse imaging and TeamViewer (Microsoft, Washington, US), embryos were monitored from a distance to allow immediate processing of embryos that showed a direct cleavage of the zygote into three or four blastomeres (multipolar division) (Movie S1). All images recorded were saved upon analysis of the cleavage kinetics (minutes) later by the Primo Vision Analyzer Software.

### Bovine genetic analysis

#### Single cell isolation and whole-genome amplification

For blastomere isolation following the first division, embryos were washed in warm TCM-199 with 10% fetal bovine serum (FBS) immediately upon the visualization of multipolar division. Thereafter, embryos were treated with pronase to dissolve the zona pellucida (0.1% protease from S. griseus, in TCM-199), and subsequently washed in TCM-199 with 10% FBS followed by Ca^+2^/Mg^+2^-free PBS with 0.05% BSA to stimulate blastomere dissociation. Next, embryos were transferred to Ca^+2^/Mg^+2-^free PBS with 0.1% polyvinylpyrrolidone (PVP) for blastomere dissociation with a STRIPPER pipet holder and a 135 μM capillary (Origio, Cooper Surgical, CT, US). When characterized by a small diameter, irregular shape and absence of a clear cell membrane a blastomere was marked as a fragment (113,114). For isolation of single cells from blastocyst-stage embryos, zygotes showing a multipolar division were cultured for 7 or 8 days, until they reached the blastocyst stage. Removal of the zona was followed by a wash step with TCM-199 with 10% FBS, a short incubation in trypsine-EDTA at 37°C, two wash steps in Ca^+2^/Mg^+2-^free PBS with 0.1% PVP and successive pipetting with a STRIPPER pipet holder and 135 μM and 75 μM capillaries (Origio, Cooper Surgical, CT, US). All blastomeres or single cells were washed in Ca^+2^/Mg^+2^-free PBS with 0.1% PVP, and transferred into a 0.2-mL PCR tube containing 2 or 4 μL of Ca^+2^/Mg^+2^-free PBS. For single cells isolated from blastocysts, a micromanipulation system was used to ensure transfer of a single-cell. Tubed blastomeres/single cells were placed immediately on dry ice and stored at −80°C until whole-genome amplification (WGA). DNA from single blastomeres/single cells and additionally, from entire blastocysts (sibling day-8) was whole-genome amplified by multiple displacement amplification (MDA) with a REPLI-g Single Cell Kit (Qiagen, Hilden, Germany) according to the manufacturer’s instructions with full or half reaction volumes for the fast 3-h protocol. The concentration of WGA DNA was determined by Qubit Broad Range Assay (Invitrogen, <u>Carlsbad, CA, US</u>A) according to the manufacturer’s protocol. Ovarian tissue from the donor cows (i.e. mothers of the respective embryos) and semen from the two bulls (i.e. fathers of the respective embryos) were used to extract bulk DNA (DNeasy Blood and Tissue kit, Qiagen, Hilden, Germany). Bulk DNA from the father and mother of one of the bulls (i.e. paternal grandparents of the respective embryos) was extracted identically from blood.

#### SNP genotyping and haplarithmisis

Whole-genome amplified products were normalized to 150 ng/μL (single cells and sibling embryos) or 50 ng/μL (bulk parental DNA) before downstream processing with the Infinium HD assay super protocol (Illumina, San Diego, CA, US). Single-cell, multi-cell sibling and bulk parental or grandparental DNA were genotyped on BovineHD BeadChips (Illumina, San Diego, California, US). Subsequently, discrete genotypes, B-allele frequency (BAF)-values and logR values were exported using Ilumina’s GenomeStudio and fed to a modified version of the siCHILD algorithm (38), siCHILD-bovine. Briefly, siCHILD-bovine is a computational workflow that deduces genome-wide (GW) single-cell haplotype and copy number profiles. For initial phasing of the parental genotypes, offspring (E01-E23) or the paternal grandparents (E24-E25) were used. Analysis of the final haplarithm plots according to siCHIld/haplarithmisis principles (38) enabled the characterization of polyploid blastomeres as either having an additional maternal or paternal copy (e.g. digynic or diandric triploidy, respectively) and blastomeres containing only paternal or maternal genomes to be labeled androgenetic or gynogenetic. Furthermore, the origin of the additional copies could be distinguished, being either mitotic or meiotic/polyspermic. A visual overview on the interpretation of the haplarithm plots can be consulted in Fig. S2.

#### Low-coverage whole-genome sequencing and interpretation

For blastomeres or fragments without haplotypes, single-cell low-coverage whole-genome sequencing was performed. Sequencing libraries were prepared starting from 500 ng for each DNA sample with the KAPA HyperPrep Kit (Hoffman-La Roche, Basel, Switzerland) according to the manufacturer’s protocol. Single-end sequencing was performed on an Illumina Hiseq 4000 device (Illumina, San Diego, CA, US). The standard Illumina primary data analysis workflow was used for base calling and quality scoring. Next, reads were demultiplexed per sample and aligned to the reference bovine genome (BosTau8). The number of raw reads in non-overlapping 10Mb bins were counted for the genomic and mitochondrial sequences and subsequently summarized in a box plot per chromosome and for the mitochondrial sequence. The mitochondrial genomes were assembled *de novo* with NOVOPlasty (115,116). The resulting assemblies were aligned against each other with MAFFT (117), followed by the construction of a phylogenetic tree with neighbor joining (118), which was visualized with Archaeopteryx (119).

### Nuclear staining

Individual blastomeres were fixed overnight in 4% paraformaldehyde and incubated in Hoechst 33342 (1:1000 dilution in PBS/PVP) for 10 min at room temperature in the dark to visualize the nuclei. Evaluation of the blastomeres nuclear content was performed the next day by fluorescent microscopy with a Leica DM 5500 B microscope with excitation filter BP 450/90 nm and a 100 W mercury lamp. Blastomeres containing one nucleus or metaphase plate were considered mononuclear, blastomeres containing no nuclear content were considered anuclear.

### Human embryo culture, biopsy and blastocyst dissociation

The development and clinical implementation of concurrent GW haplotyping and copy number profiling of single blastomeres during preimplantation genetic testing (PGT) (38,93,94) provides a screen of genome-wide error profiles in human embryos fertilized through ICSI. In a retrospective study, 2300 day-3 single blastomere biopsies derived from 2257 cleavage-stage embryos which reached the blastocyst-stage were analysed (96). Embryo culture, biopsy and biopsy processing for these embryos was performed under a standard clinical workflow for PGT at UZ Leuven, as previously described by (120). GW ploidy violations, such as haploidy/GW UPD and triploidy, were detected in 2.4%. These embryos are deemed not eligible for transfer. We classified those embryos as gynogenetic (carrying only maternal DNA) or androgenetic (carrying only paternal DNA), and triploid embryos as digynic or diandric, respectively. Ten cleavage-stage embryos with either an androgenetic (n=2) or a gynogenetic (n=8) blastomere were followed up by thawing and dissociation of the blastocyst for single-cell analysis. Manipulations of the whole blastocyst were performed with a STRIPPER pipette with 175 or 135 μm capillaries (Origio, CooperSurgical, Inc., USA). The dissociation procedure was performed as follows: a short incubation of the blastocyst in Acidic Tyrode’s solution (Sigma-Aldrich, Merck KGaA, Germany) was executed until visual disappearance of the zona pellucida was observed. The blastocyst was consecutively washed in three drops of biopsy medium (LG PGD Biopsy Medium, Life Global) and incubated in trypsin at 37 °C. Subsequently, the blastocyst was washed three times in biopsy medium. Individual cells from the blastocysts were then isolated by manual pipetting using a STRIPPER pipette with a 75 μm capillary (Origio, CooperSurgical, Inc., USA) and washed three times 1% PVP-PBS. Subsequently, each isolated cell was transferred into a 0.2 ml PCR tube with 2 μl PBS and stored at -20°C until further use. Samples were then whole genome amplified using REPLI-g Single Cell Kit (Qiagen, Germany) for half-volume reactions and with incubation at 30°C for 2 h followed by 65°C during 10 min for inactivation. Analysis was performed using siCHILD/haplarithmisis (38), as mentioned previously under the standard clinical workflow for PGT at UZ Leuven. For both human blastocysts, E01 and E02, an (overall) biparental balanced single blastomere from a sibling embryo (for E02: a maternal meiotic trisomy 15 was present) was used to execute phasing of the blastocysts’ single cells.

### Statistical analyses and data visualization

Statistical analyses were conducted using R (Version 3.6.1.). Linear mixed models were built to determine the effect of the cleavage pattern on the time of cleavage. The family to which the embryo belonged to and the embryo from which the blastomere originated were included as random effects. Statistical significance level was set at P≤0.05. Circos plots were drawn using the Circlize package in R (121).

## Declarations

### Ethics approval and consent to participate

Ethical review was waived for bovine embryonic samples as they were produced from slaughterhouse-derived materials and are no subject of ethical approval. Ethical approval for the human embryonic samples was granted by Ethical Committee of UZ/KU Leuven (S59351). All patients received information on the study and provided informed consent on the use of their data.

### Consent for publication

Not applicable.

### Availability of data and materials

Human embryo, parental and phasing relatives’ raw genotyping data has been deposited at the European Genome-phenome Archive (EGA), which is hosted by the EBI and the CRG, under accession number EGAS00001005543. It is available to academic users upon request to the Data Access Committee (DAC) of KU Leuven via the corresponding author (J.R.V). The bovine sequencing data for this study have been deposited in the European Nucleotide Archive (ENA) at EMBL-EBI under accession number PRJEB46925 (https://www.ebi.ac.uk/ena/browser/view/PRJEB46925). The bovine SNP array genotyping data discussed in this publication have been deposited in NCBI’s Gene Expression Omnibus and are accessible through GEO Series accession number GSE182345 (https://www.ncbi.nlm.nih.gov/geo/query/acc.cgi?acc=GSE182345). In compliance with the GDPR (General Data Protection Regulation 2016/679) and the study protocol, the human PGT-M blastocyst dataset used in the study is not publicly available.

### Author contributions

O.T., K.S., A.V.S. and J.R.V. conceived and designed the experiments. S.D. and K.P. led patient counselling, IVF– PGT associated procedures. and initial human PGT sample analysis. T.D.C. and M.C. performed the bovine experiments. H.M. performed human blastocyst experiments, human and bovine analysis for genome-wide haplotyping and copy number profiling and sequence alignment. T.D.C., H.M. and O.T. assisted with SNP-array technology. N.D. performed the *de novo* mitochondrial genome assembly. T.D.C., H.M., O.T. and J.R.V. performed data analysis. T.D.C. and J.R.V. wrote the original draft of the manuscript. T.D.C, H.M., O.T., N.D., K.S., A.V.S., and J.R.V. edited the final draft of the manuscript. K.S., A.V.S. and J.R.V. supervised the study.

## Acknowledgments

The authors are grateful to the Genomics Core for facilitating the use of sequencing technologies, to P. Van Damme for her excellent technical assistance, to the Euro Meat Group for their help in the collection of bovine ovaries and to CRV for providing the bovine sperm. We thank dr. Aspasia Destouni and Kate E. Stanley for a critical reading of the manuscript.

## Additional Files

### Additional Figures S1, S2, S3 (Additional Figures.pdf)

Fig. S1. **Anuclear blastomeres and fragments. A)** Phylogenetic tree based on reassembled mitochondrial genomes of 11 anuclear blastomeres and two anuclear fragments, showing a common ancestor for sequenced blastomeres and fragments retrieved from the same embryo. One anuclear blastomere was not included. **B)** Box plots and median values of the number of raw reads per 10 Mb bin (y-axis) per chromosome (chr1-chrX) and for the mitochondrial DNA (chr M) (x-axis) as determined by single-cell low-coverage whole-genome sequencing for 12 anuclear blastomeres and two anuclear fragments demonstrate the abundance of mitochondrial DNA. **C)** Identical plots as in B, excluding chr M demonstrate the presence of fragments of chromosomal fragments in three anuclear blastomeres. **D)** Overlay of bright field and Hoechst fluorescent image show anuclear (1) and mononucleated (2;3) blastomeres resulting from a multipolar zygotic division in three blastomeres.

Fig. S2. **Analysis of blastomeres following multipolar zygotic division A)** Interpretation of haplarithm plots. Overview of chromosome-wise haplarithm patterns for distinct genomic constitutions (i.e. normal disomy, paternal monosomy and paternal meiotic/dispermic uniparental heterodisomy). Corresponding whole-genome errors (i.e. biparental diploid or androgenetic) are characterized by the manifestation of those patterns throughout the (majority of the) genome. Defined single-cell BAF-values of the segmented P1, P2, M1 and M2, form haplotype blocks, demarcated by pairwise breakpoints, i.e., homologous recombinations. Haplotype blocks, as well as the distance between the P1-P2 or M1-M2 in the paternal and maternal haplarithm, respectively, and the positioning of homologous recombinations, denote the origin and nature of copy number. The normalized logR-values are integrated with haplarithm profiles for copy number profiling. Principles of interpretation are according to Zamani Esteki et al. (2015) (38). **B)** An overview of haplarithm profiles of 82 blastomeres and two fragments (grey squares) is depicted per category of whole-genome segregation profiles, as discussed in the main text. Each embryo is identified by a description at the top left of the embryo ID and cross (EmbryoID_Embryocross). At the top right, three chronological time-lapse images of the cleaving zygote are depicted. From left to right, the pictures show the initiation of the cleavage furrow, the ongoing first division and the embryo immediately after cleavage and before cell isolation (when video available). For each embryo, a schematic representation of likely steps leading to the genomic profile of each blastomere (B1-B4) or fragment (F1) is given. Chromosome-wise interpretation (1 - X) per blastomere is visualized in the bar above the haplarithm plots (see legend). Below each bar, the paternal haplarithm (pat-BAF), the maternal haplarithm (mat-BAF) and the normalized LogR-values (LogR) are depicted. Paternal cross-over sites are depicted by the arrows (black, green or blue). A combination of parental cross-over sites in one blastomere or different cross-over sites in blastomeres of the same embryo uncover polyspermic fertilization or a meiotic error. Maternal cross-over sites (red, pink, orange) were only depicted in gynogenetic blastomeres and in case of whole-genome meiotic errors.

Fig. S3. **Genome-wide composition of blastocysts following multipolar zygotic division**. Circos plots of six embryos that developed to the blastocyst-stage following multipolar zygotic division in which each circle represents the interpreted genome constitution per chromosome (1 - X) of a sampled single cell. The interpreted genome spanning the largest part of the chromosome was chosen as overall interpretation per chromosome, as such, segmental chromosomal errors are not depicted. On the left of each circus plot, a picture is shown of each embryo before dissociation (not available for E26_Cross13).

### Additional movie S1 (Additional Movie.mp4)

Movie S1: **bipolar and multipolar zygotic division**. Movie showing a bipolar zygotic division and a multipolar zygotic division in three or four cells, respectively.

