## Additional figures S1,S2 and S3 for "Parental genomes segregate into different blastomeres during multipolar zygotic divisions leading to mixoploid and chimeric blastocysts"

### Supplemental Figures

#### Figure S1

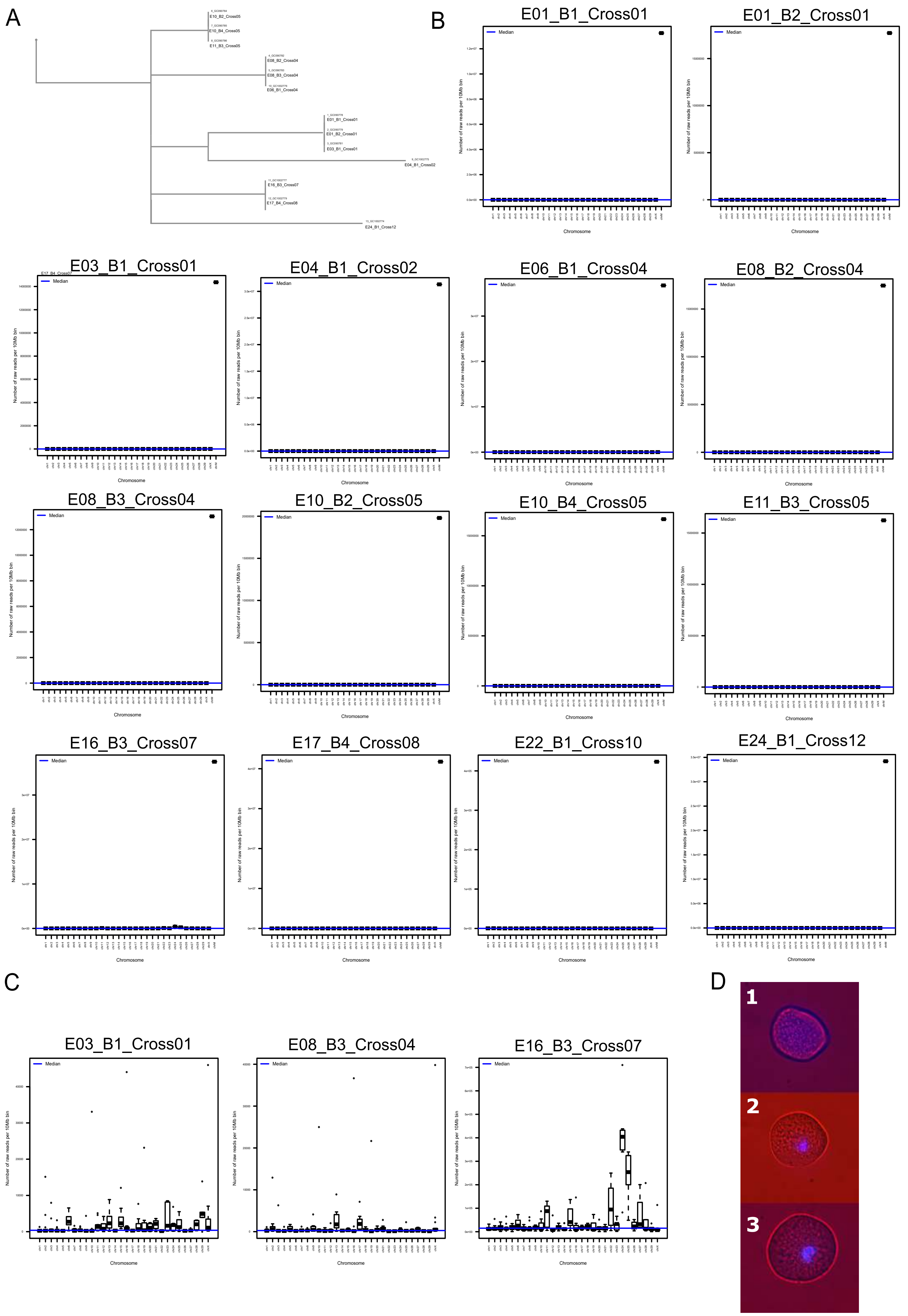

Figure S2

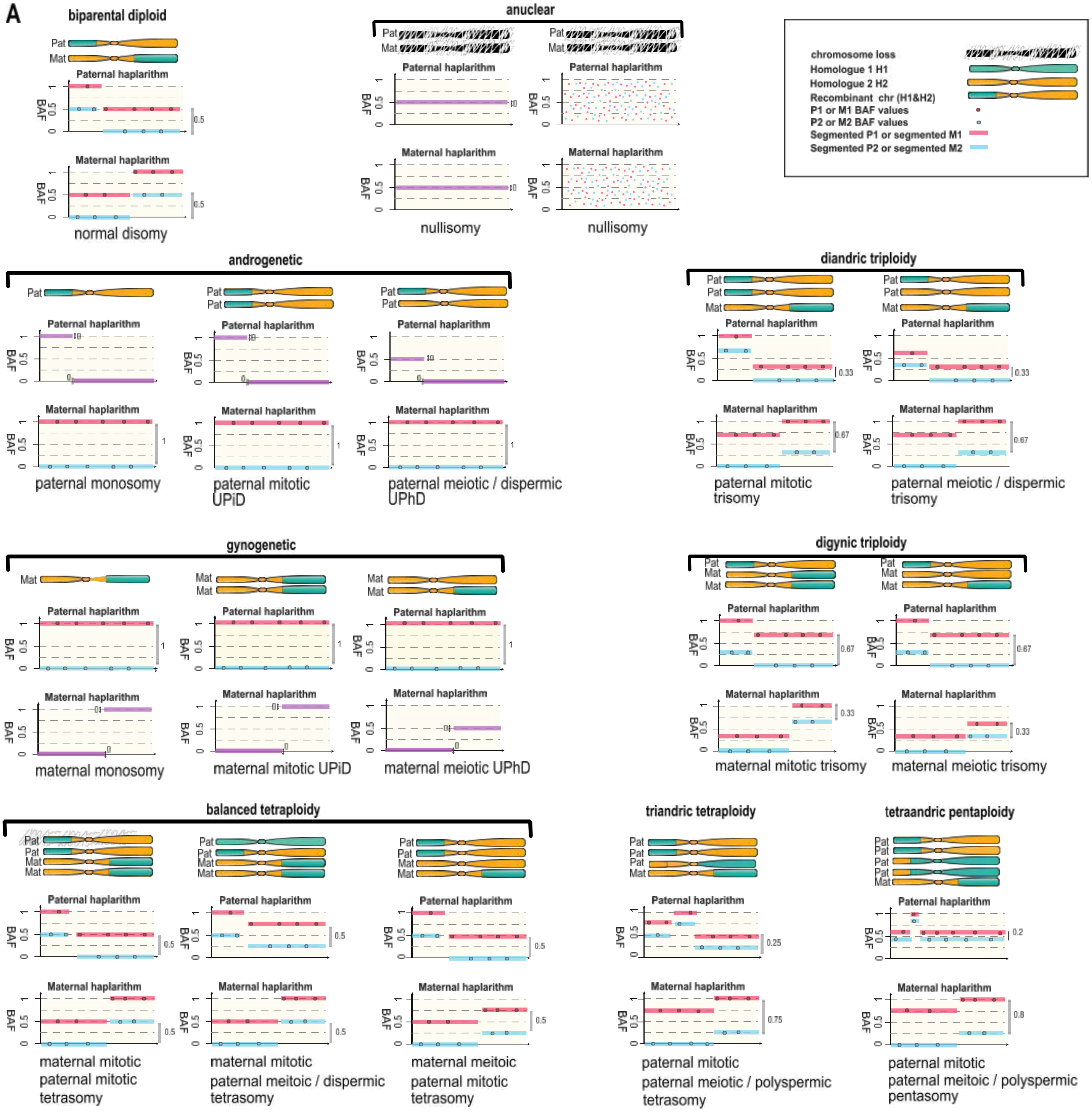
A
biparental diploid
Pat
Mat
Paternal haplarithm
Maternal haplarithm
BAF
normal disomy
anuclear
nullisomy
nullisomy
chromosome loss
Homologue 1 H1
Homologue 2 H2
Recombinant chr (H1&H2)
P1 or M1 BAF values
P2 or M2 BAF values
Segmented P1 or segmented M1
Segmented P2 or segmented M2
androgenetic
paternal monosomy
paternal mitotic UPiD
paternal meiotic / dispermic UPhD
diandric triploidy
paternal mitotic trisomy
paternal meiotic / dispermic trisomy
gynogenetic
maternal monosomy
maternal mitotic UPiD
maternal meiotic UPhD
digynic triploidy
maternal mitotic trisomy
maternal meiotic trisomy
balanced tetraploidy
maternal mitotic paternal mitotic tetrasomy
maternal mitotic paternal meiotic / dispermic tetrasomy
maternal meitoic paternal mitotic tetrasomy
triandric tetraploidy
paternal mitotic paternal meiotic / polyspermic tetrasomy
tetraandric pentaploidy
paternal mitotic paternal meiotic / polyspermic pentasomy

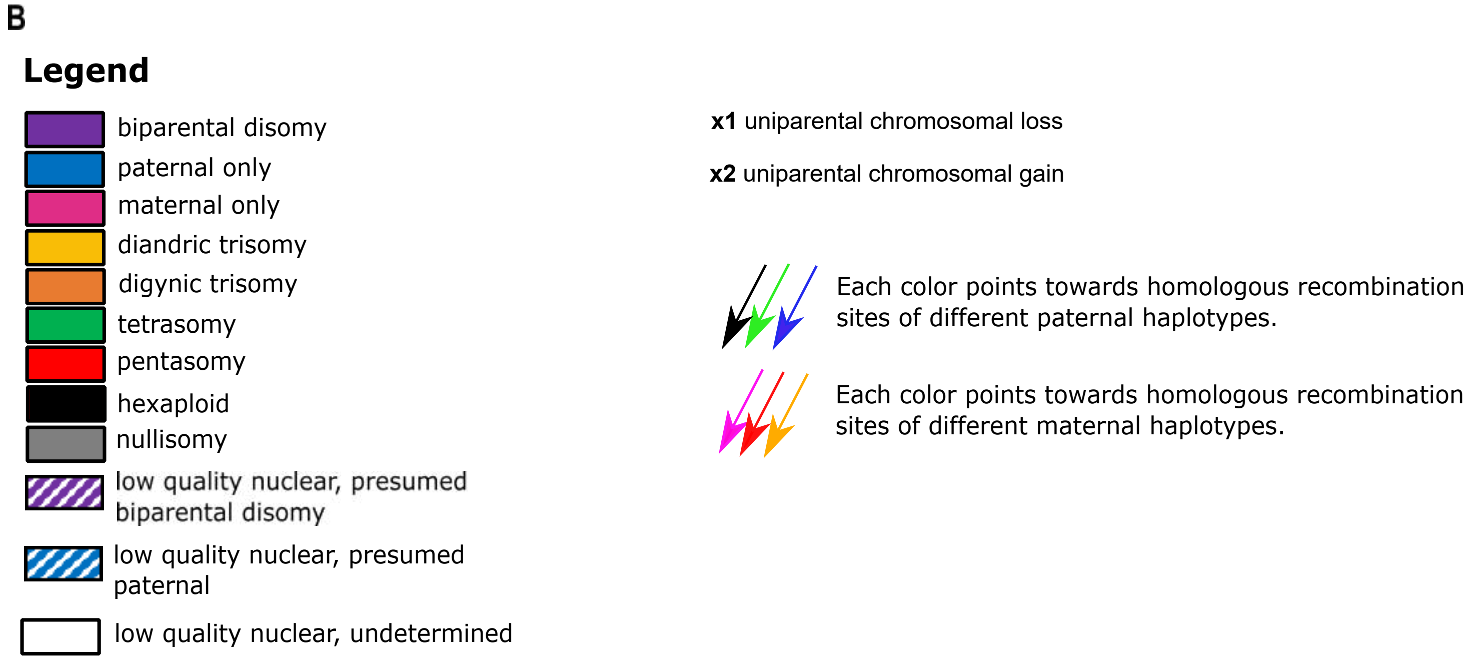
B
Legend
biparental disomy
paternal only
maternal only
diandric trisomy
digynic trisomy
tetrasomy
pentasomy
hexaploid
nullisomy
low quality nuclear, presumed biparental disomy
low quality nuclear, presumed paternal
low quality nuclear, undetermined
x1 uniparental chromosomal loss
x2 uniparental chromosomal gain
Each color points towards homologous recombination sites of different paternal haplotypes.
Each color points towards homologous recombination sites of different maternal haplotypes.

Figure S2B (continued)

**1. Embryos consisting of diandric triploid, biparental diploid and androgenetic blastomeres**

E05_Cross03

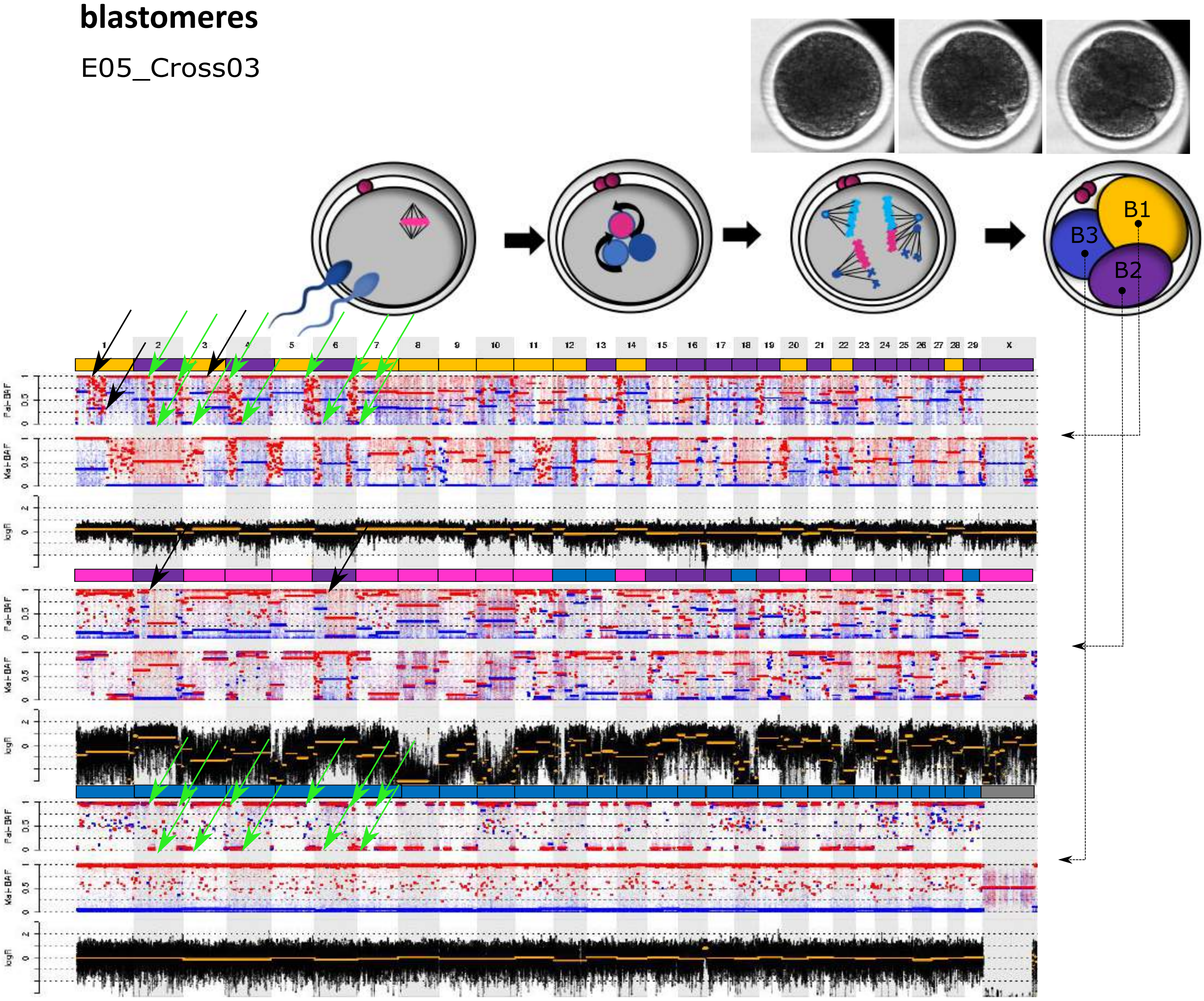

Figure S2B (continued)

E07_Cross04

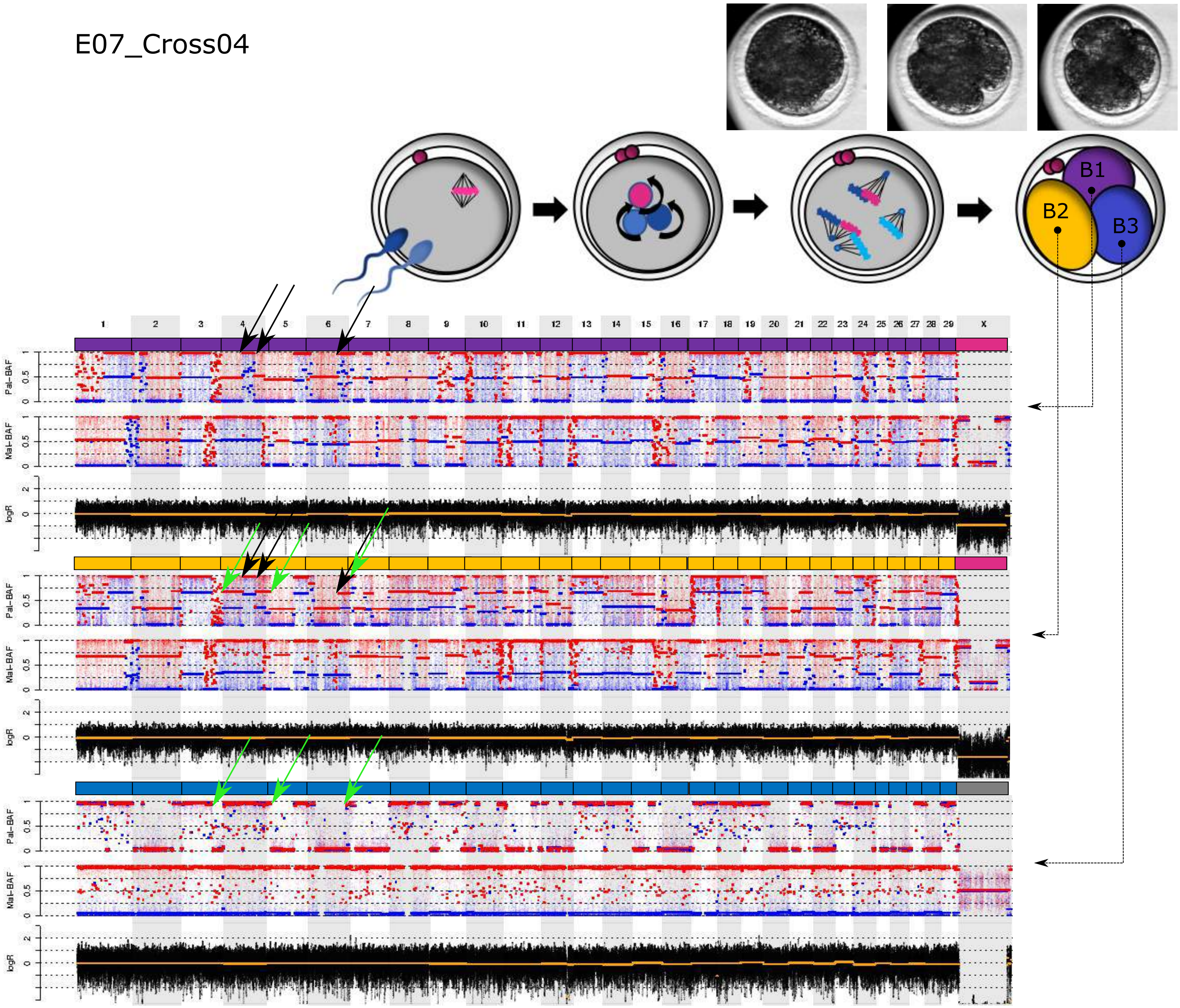

Figure S2B (continued)

E09_Cross04

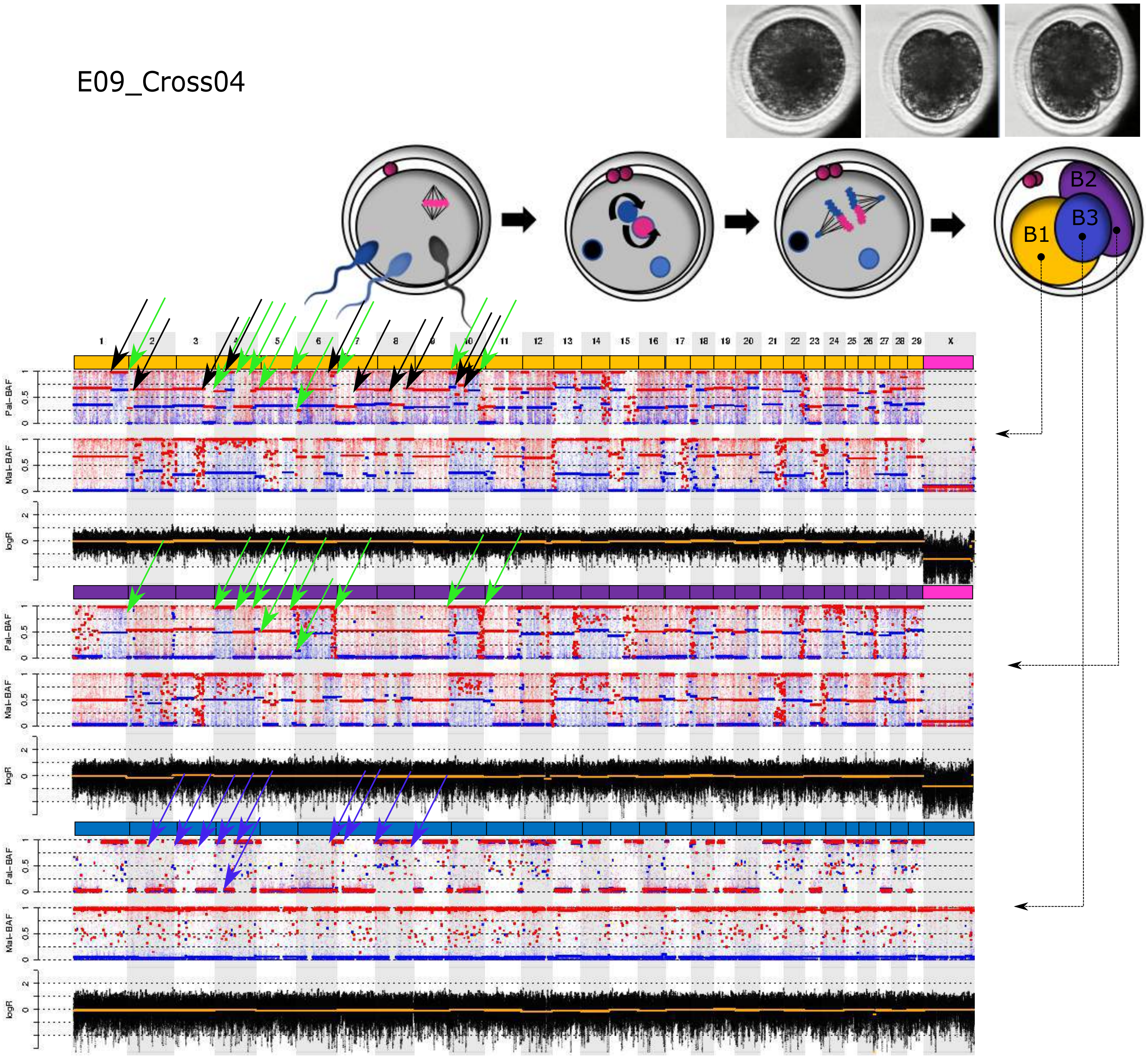

Figure S2B (continued)

### 2. Embryos consisting of biparental and androgenetic blastomeres

E02_Cross01

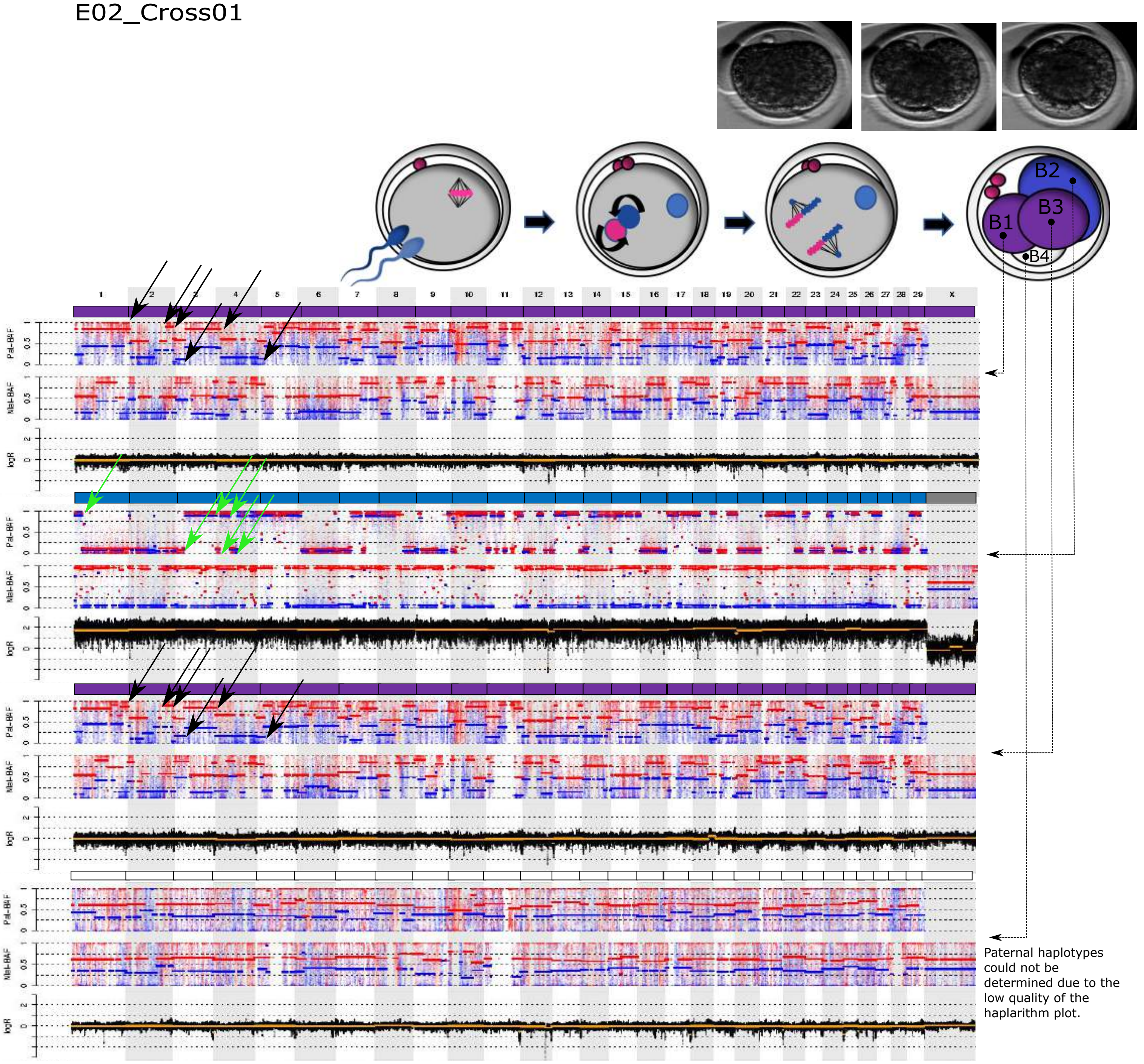

Figure S2B (continued)

E06_Cross04

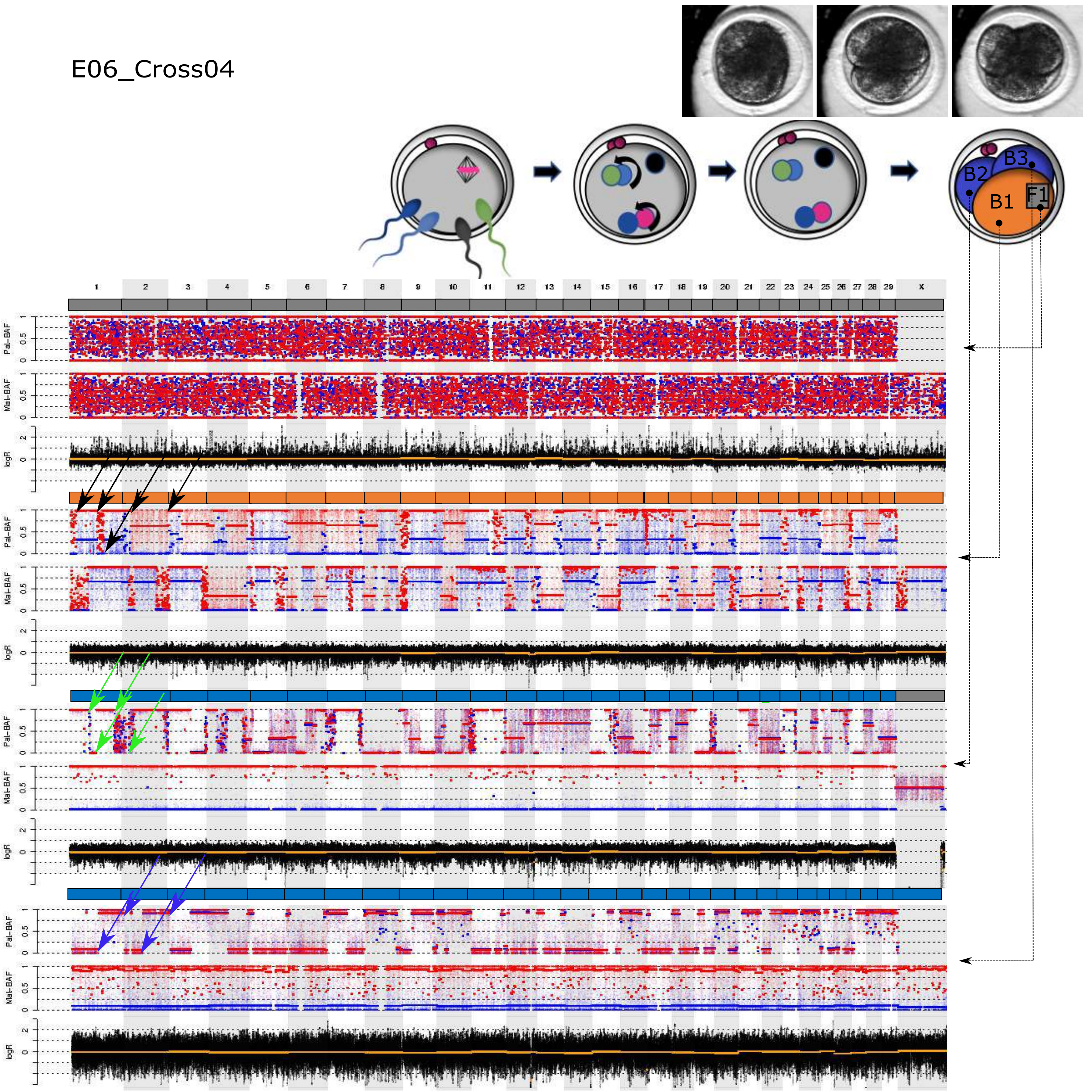

Figure S2B (continued)

E11_Cross05

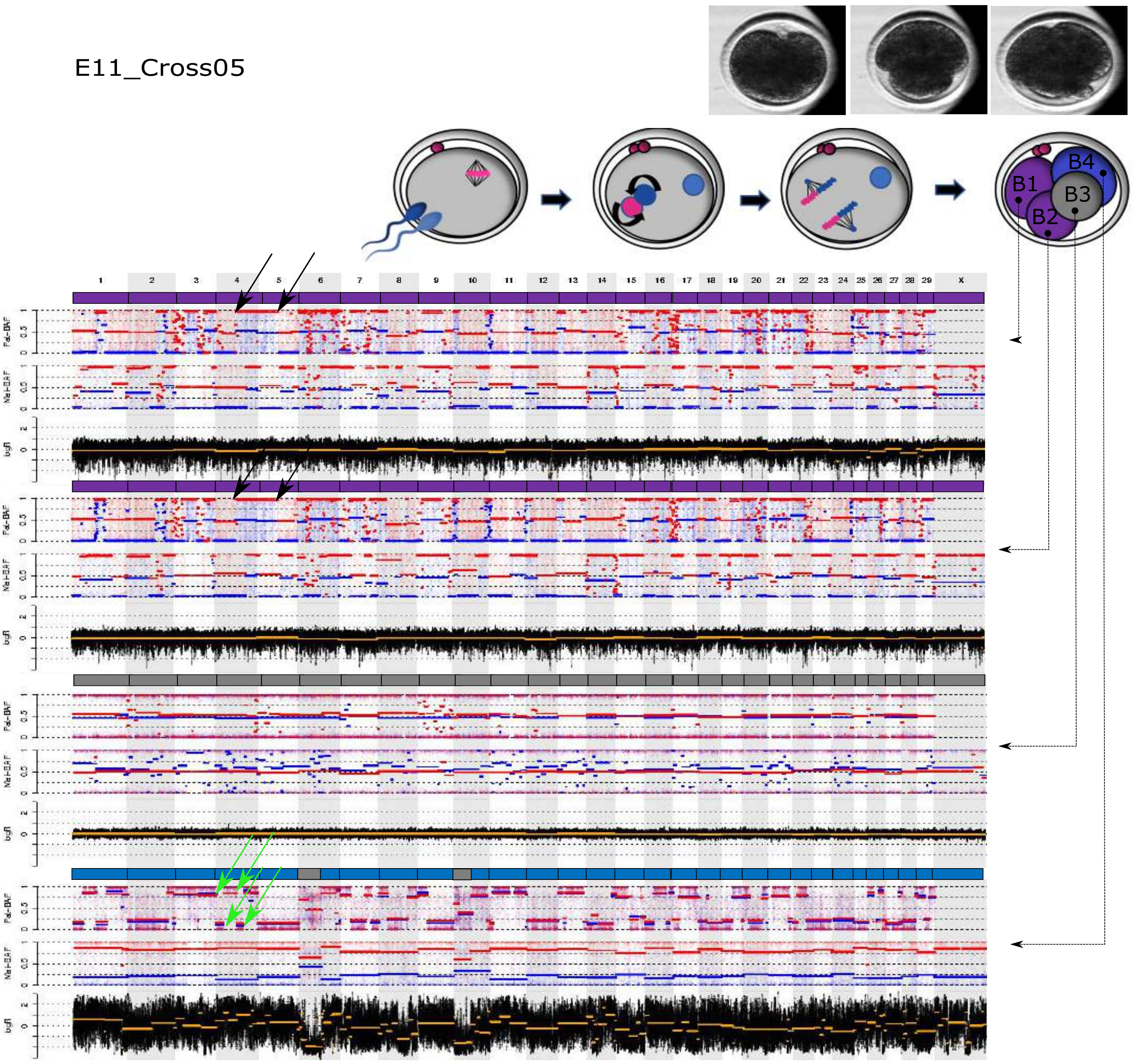

Figure S2B (continued)

E12_Cross05

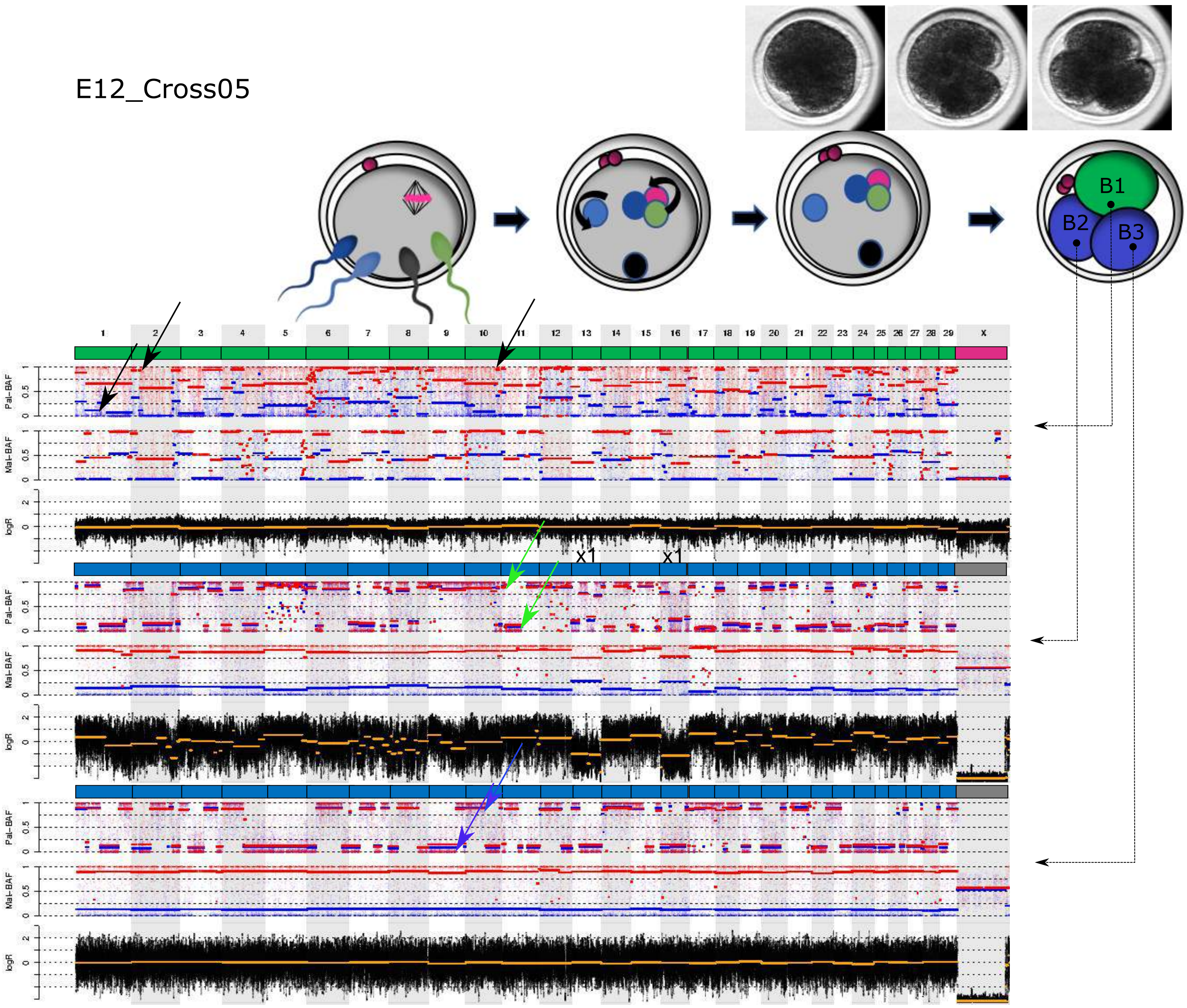

Figure S2B (continued)

E13_Cross11

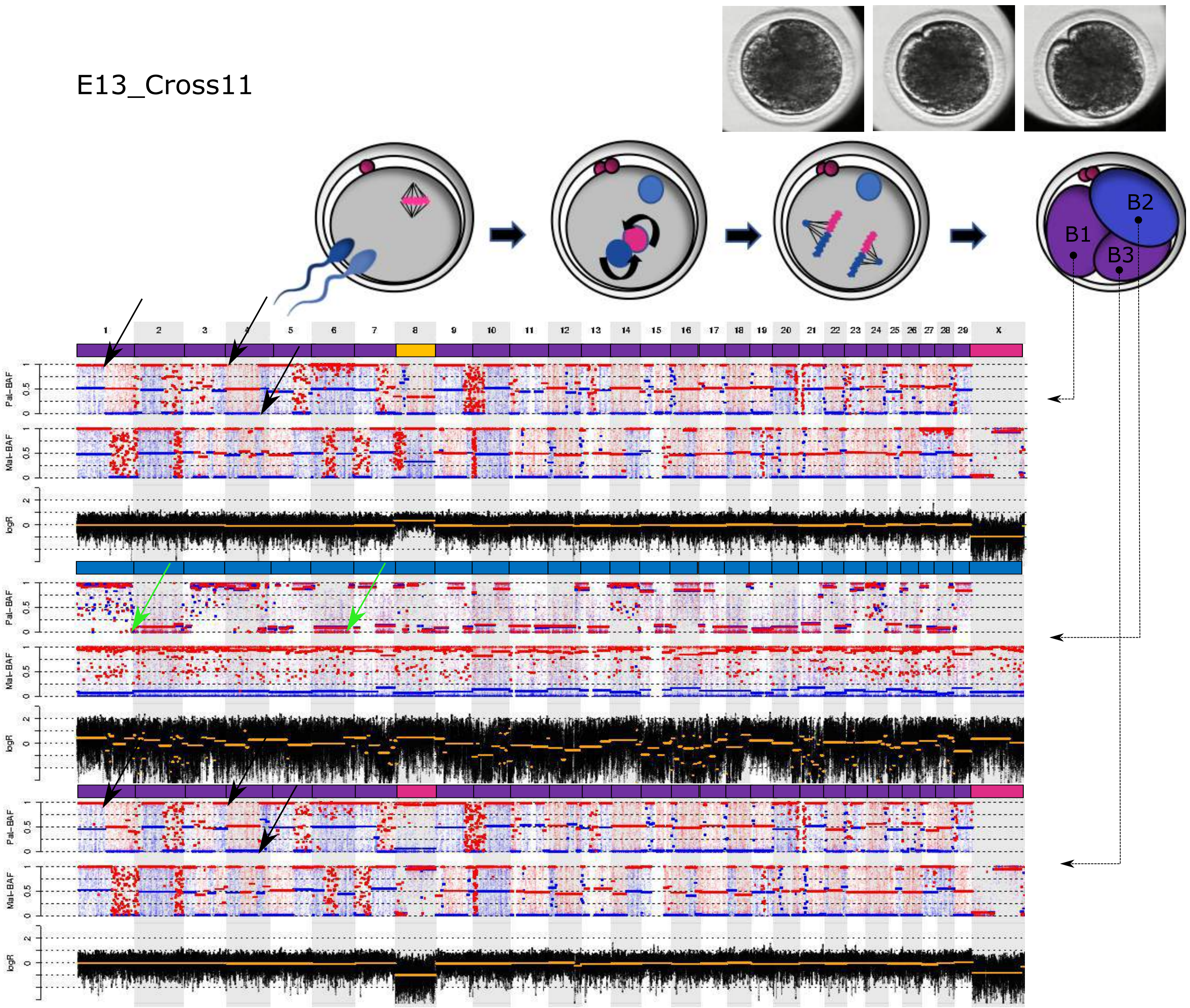

Figure S2B (continued)

E14_Cross07

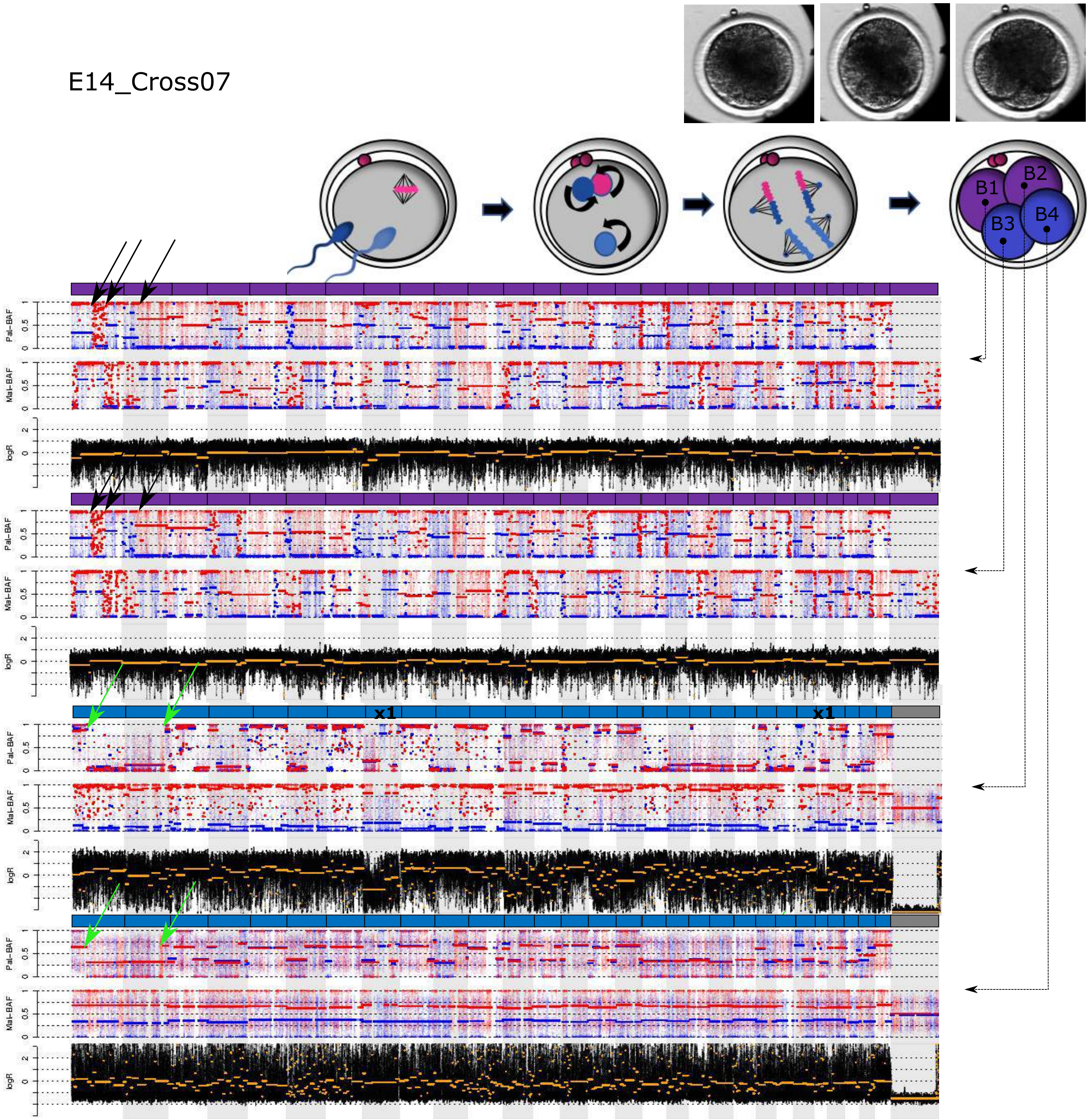

Figure S2B (continued)

E15_Cross07

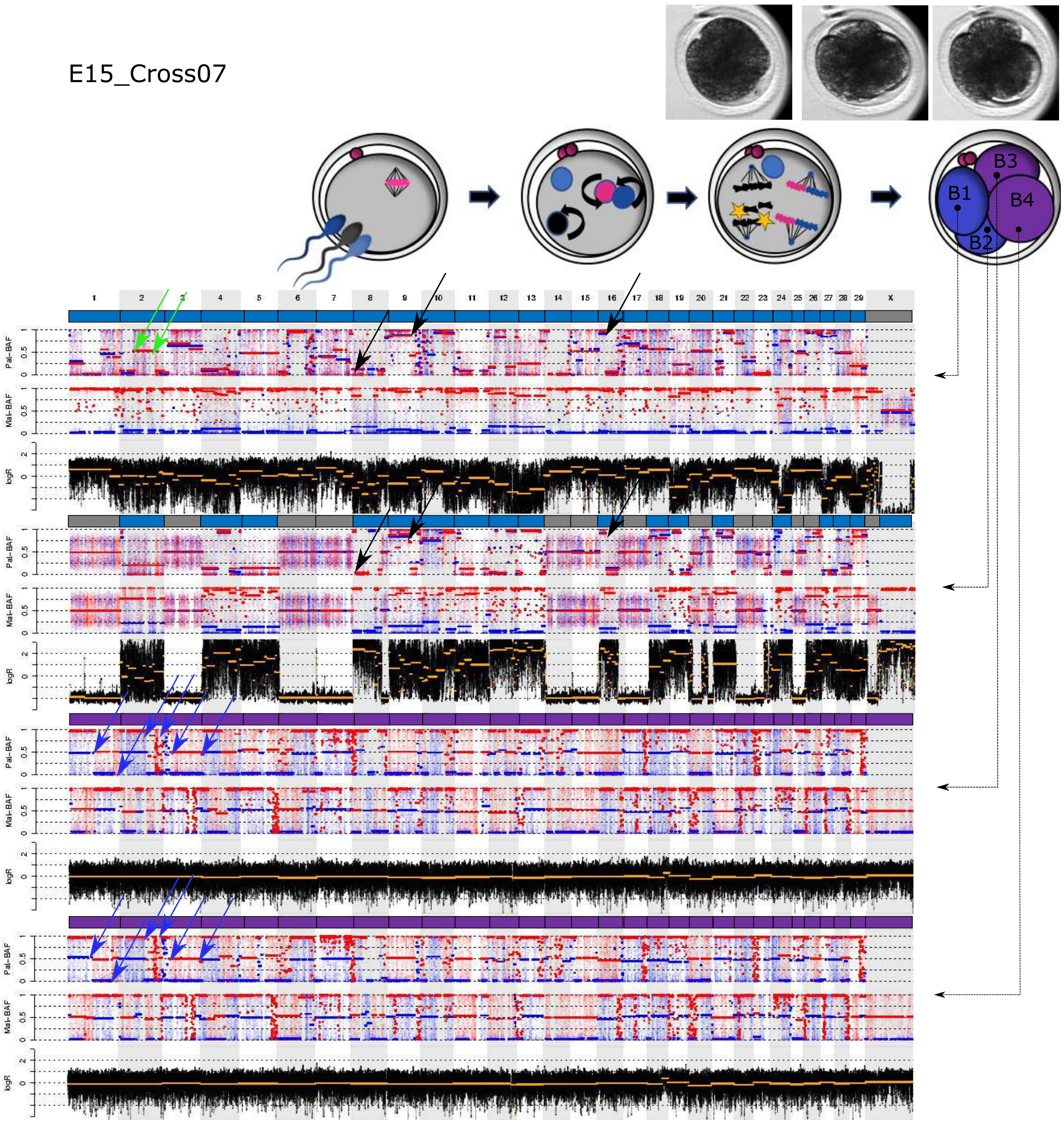

Figure S2B (continued)

E16_Cross07

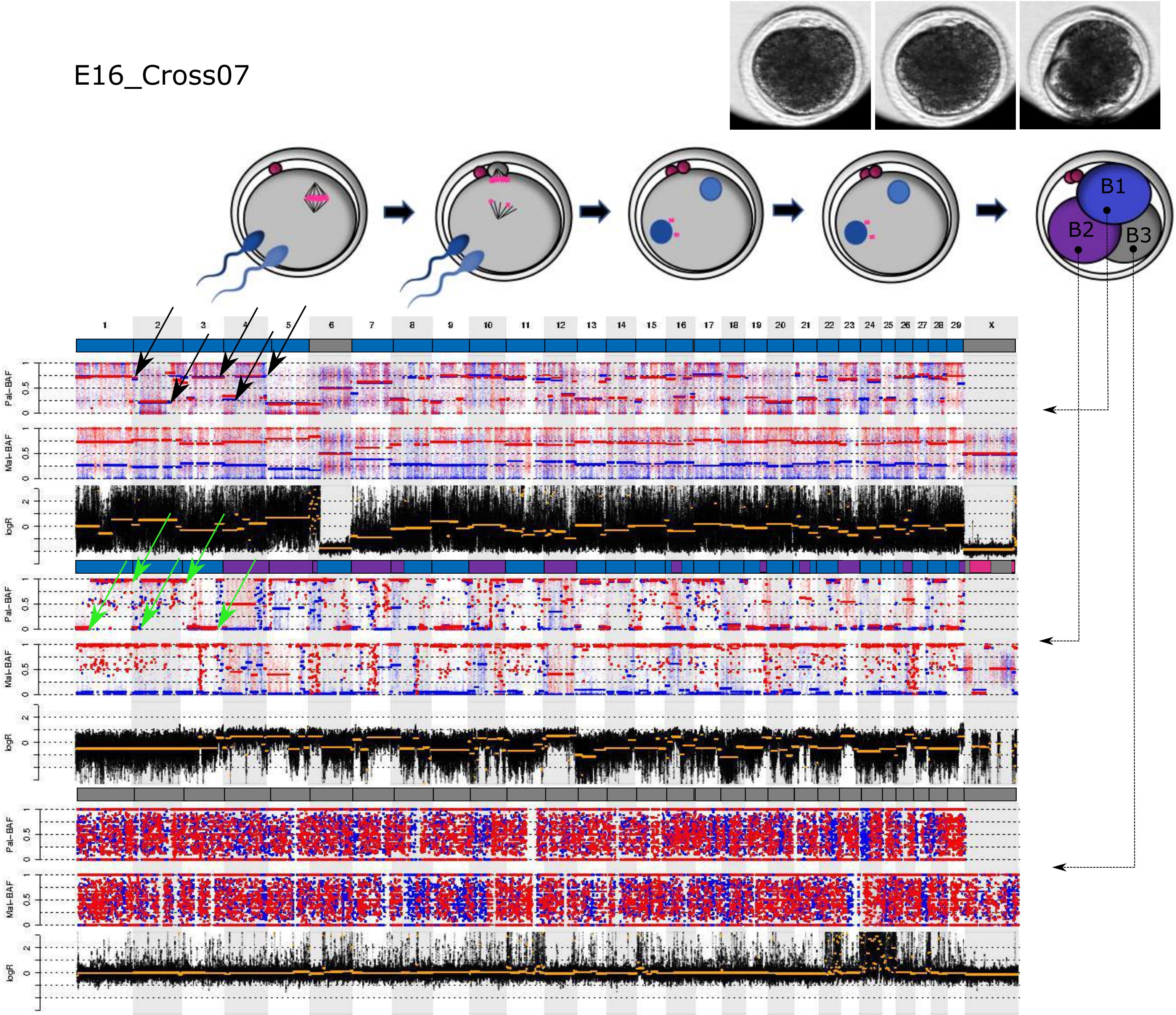

Figure S2B (continued)

E17_Cross07

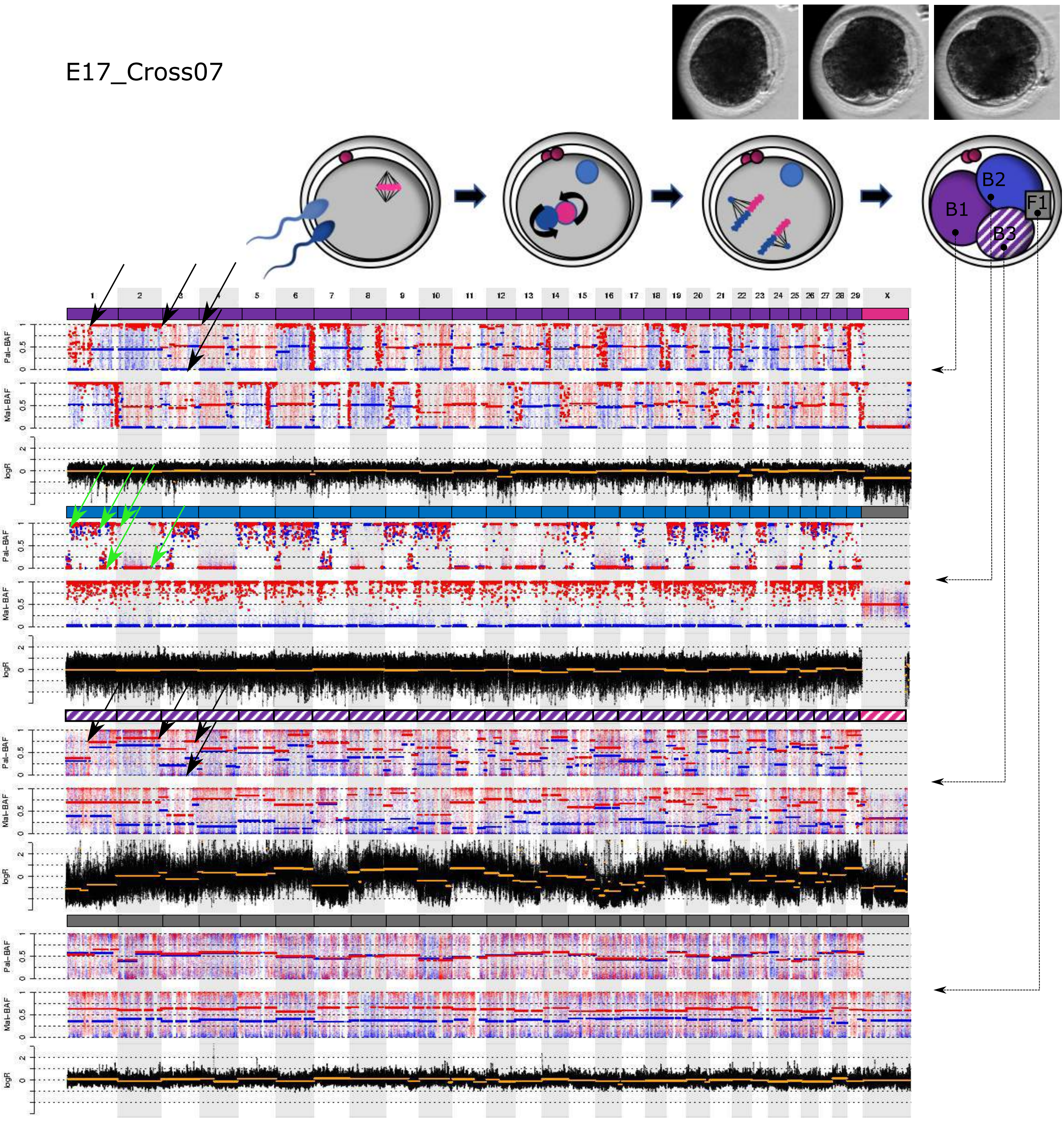

Figure S2B (continued)

E18_Cross08

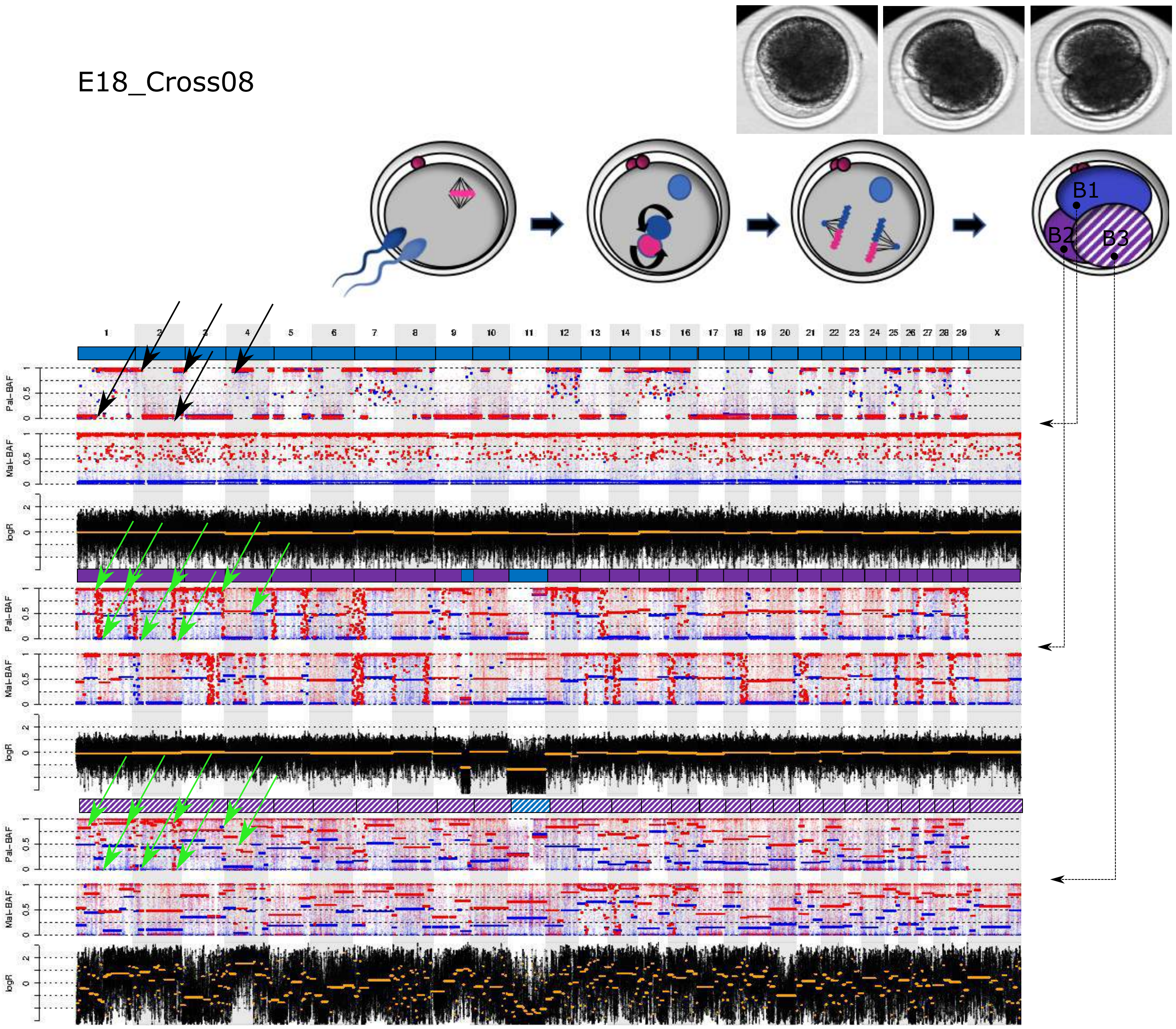

Figure S2B (continued)

E19_Cross08

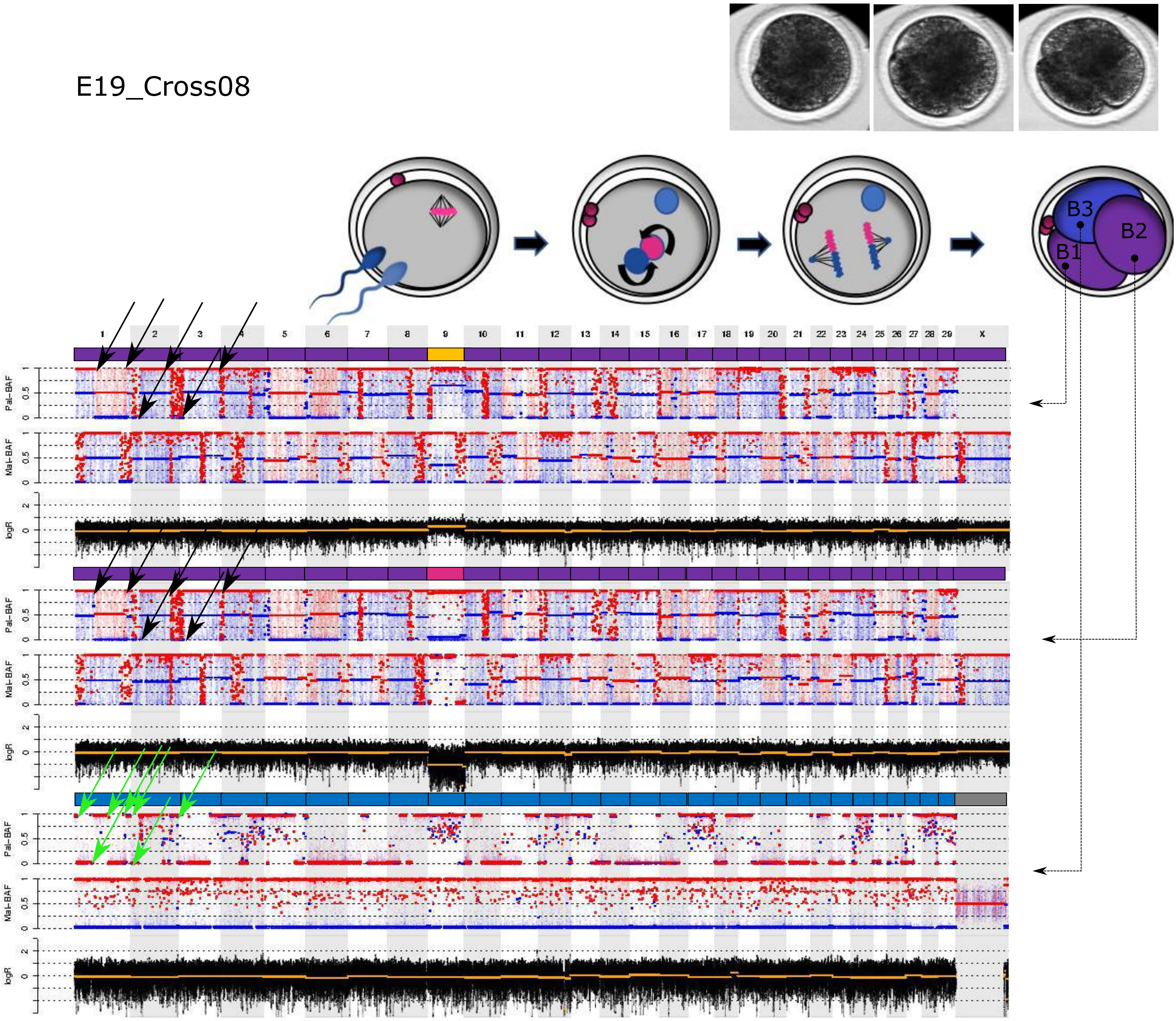

Figure S2B (continued)

E21_Cross09

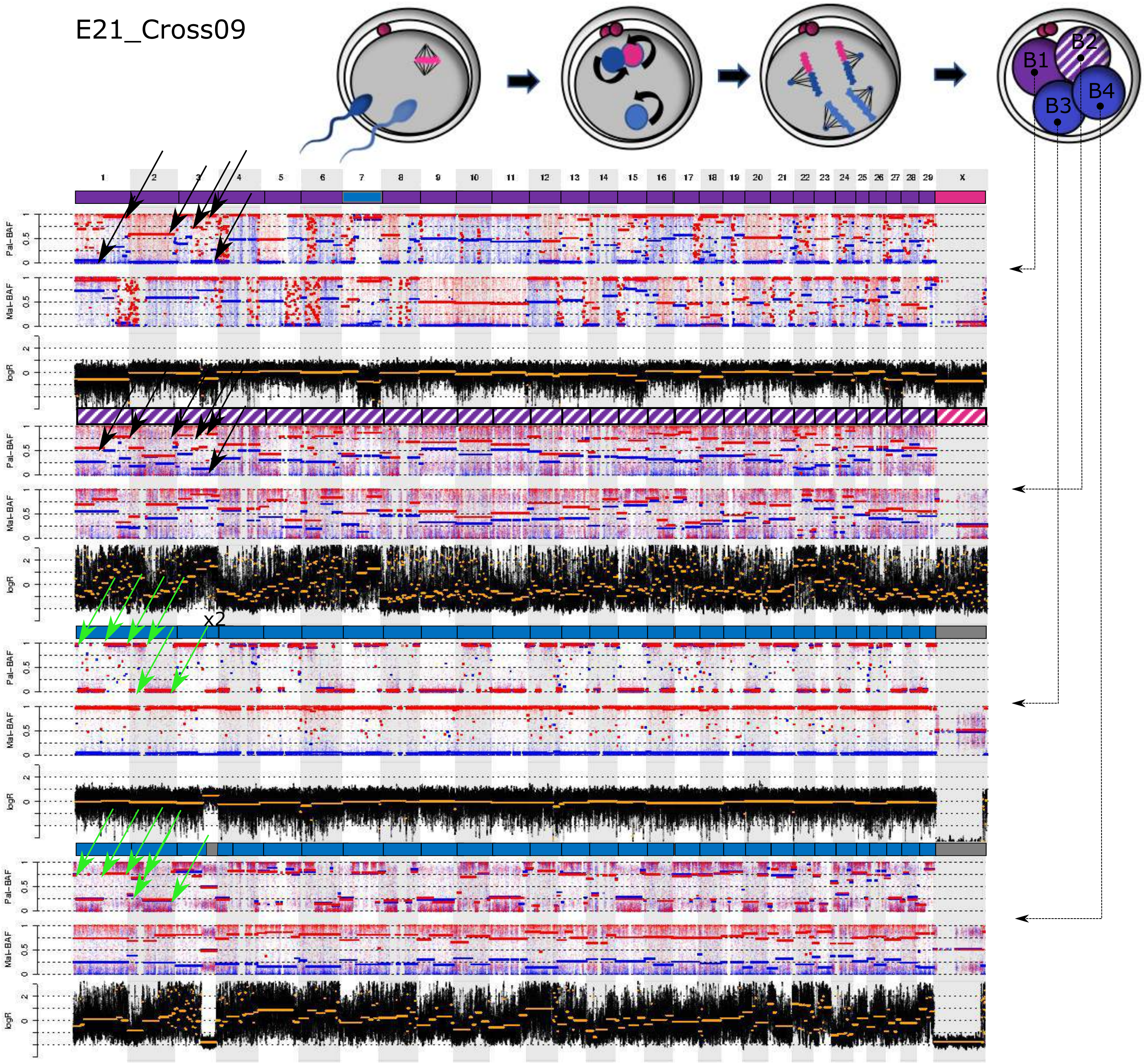

Figure S2B (continued)

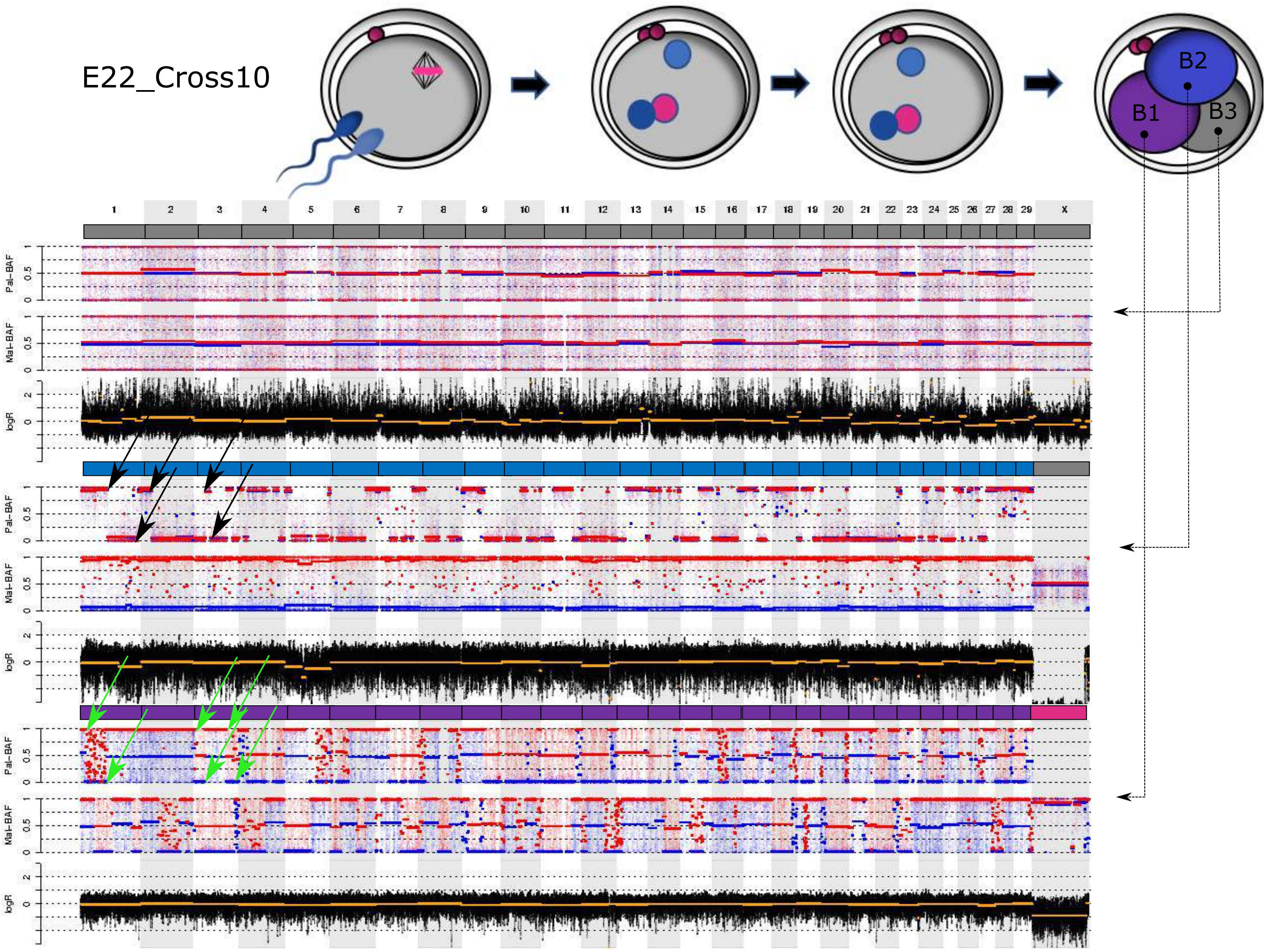

Figure S2B (continued)

### 3. Embryos consisting of androgenetic and gynogenetic blastomeres

E23_Cross11

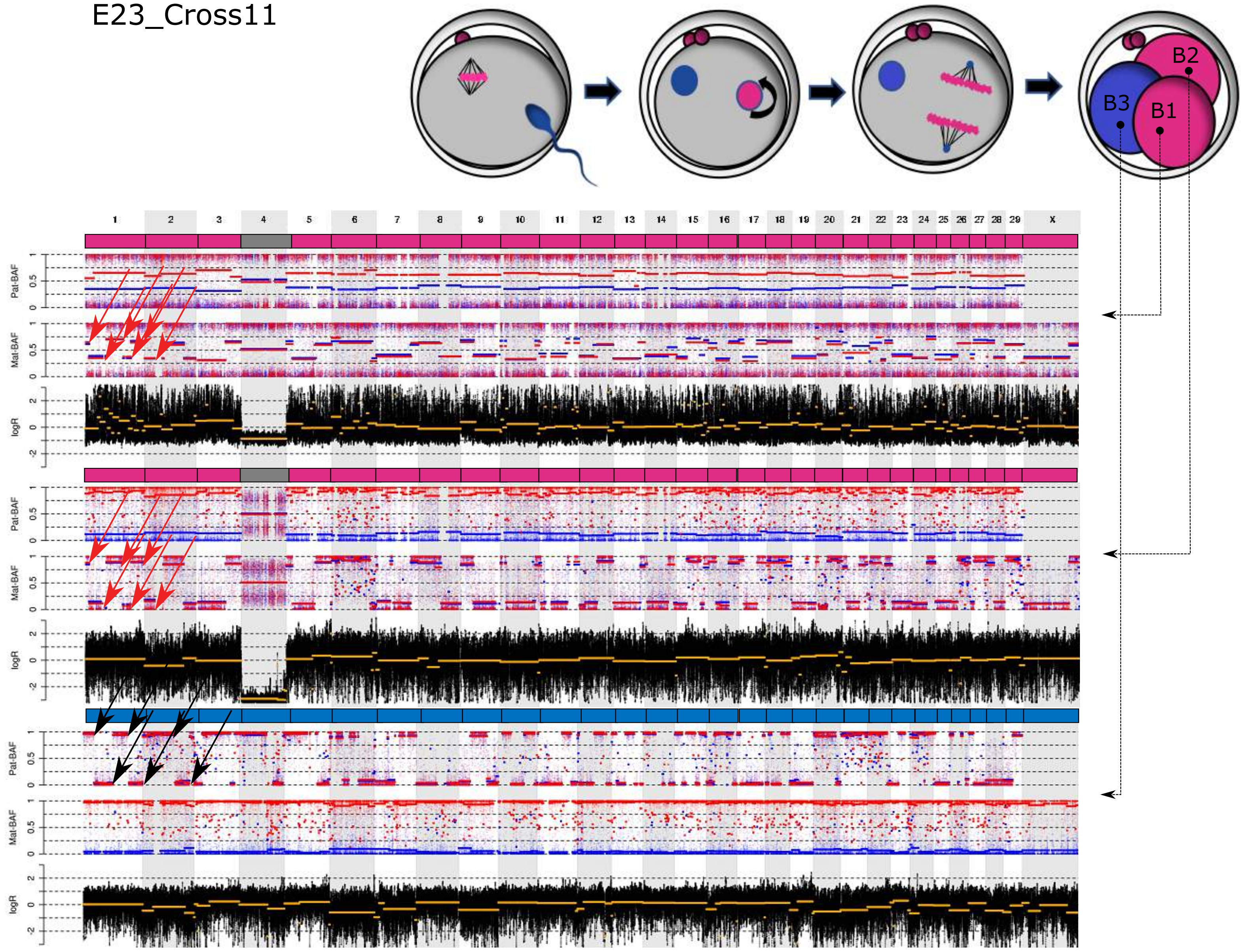

Figure S2B (continued)

E25_Cross12

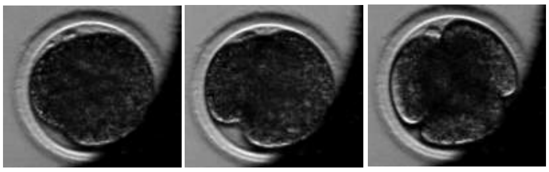

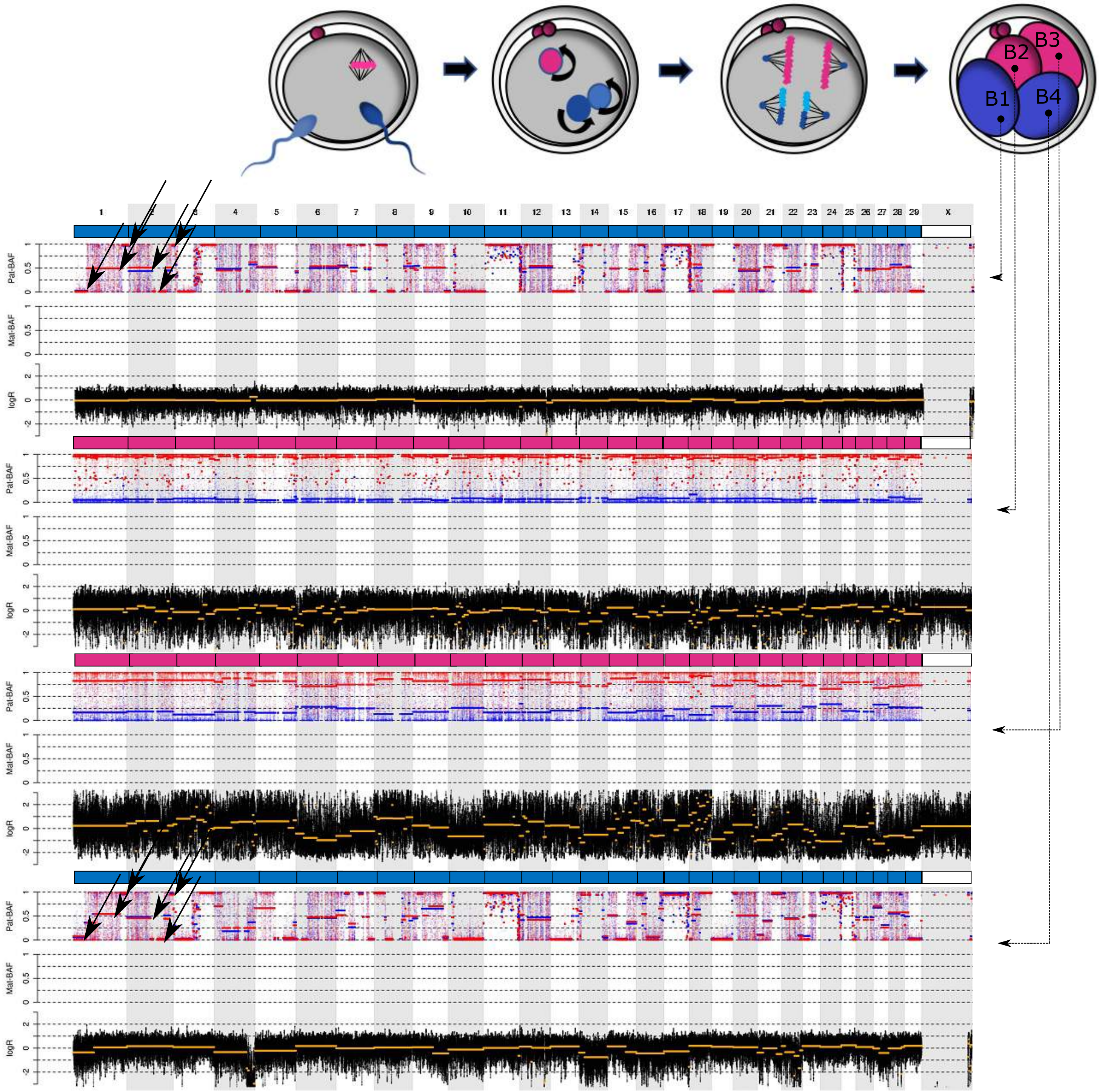

Figure S2B (continued)

### 4. Androgenetic embryos

E01_Cross01

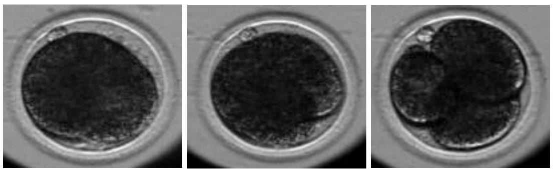

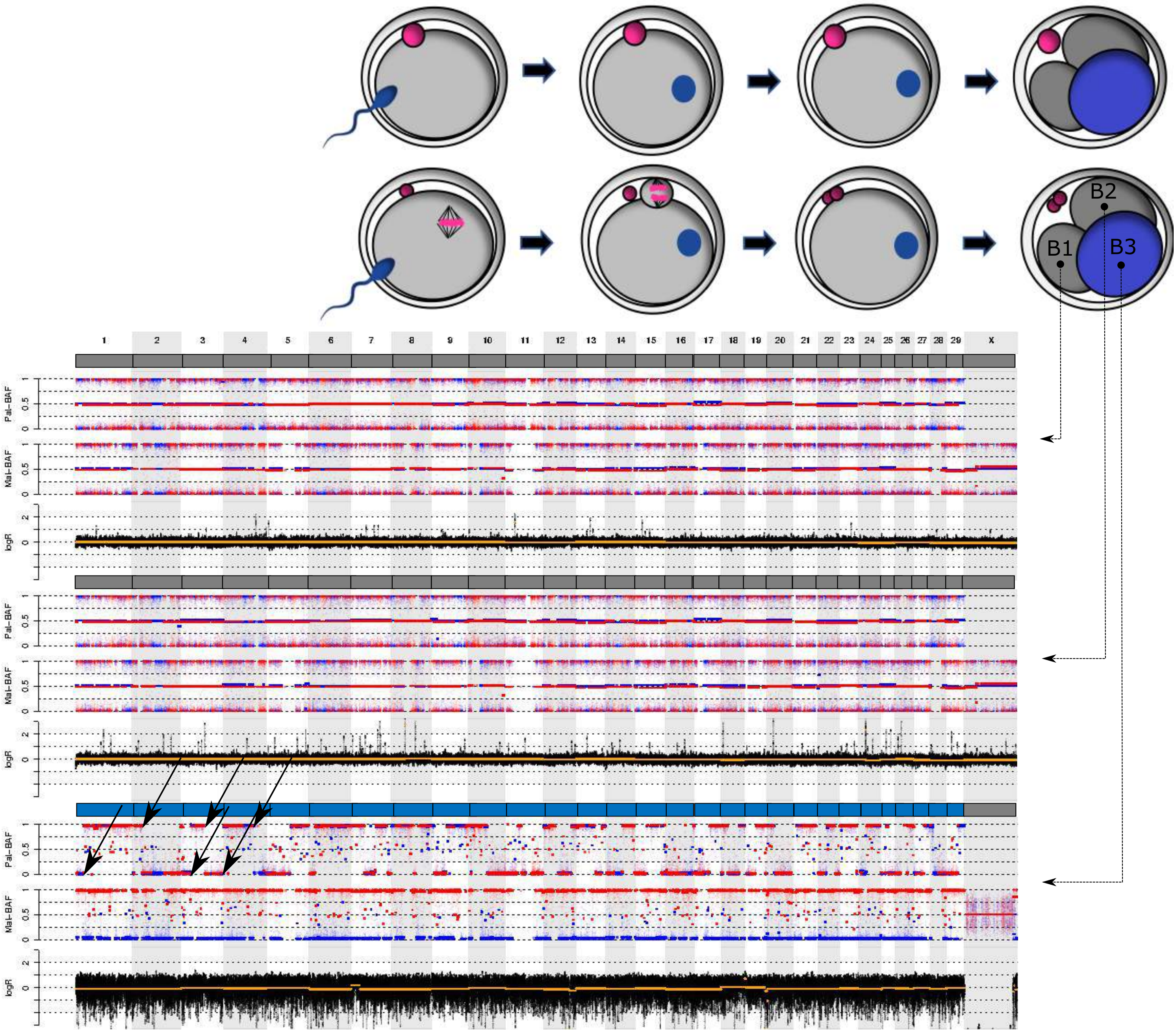

Figure S2B (continued)

E10_Cross05

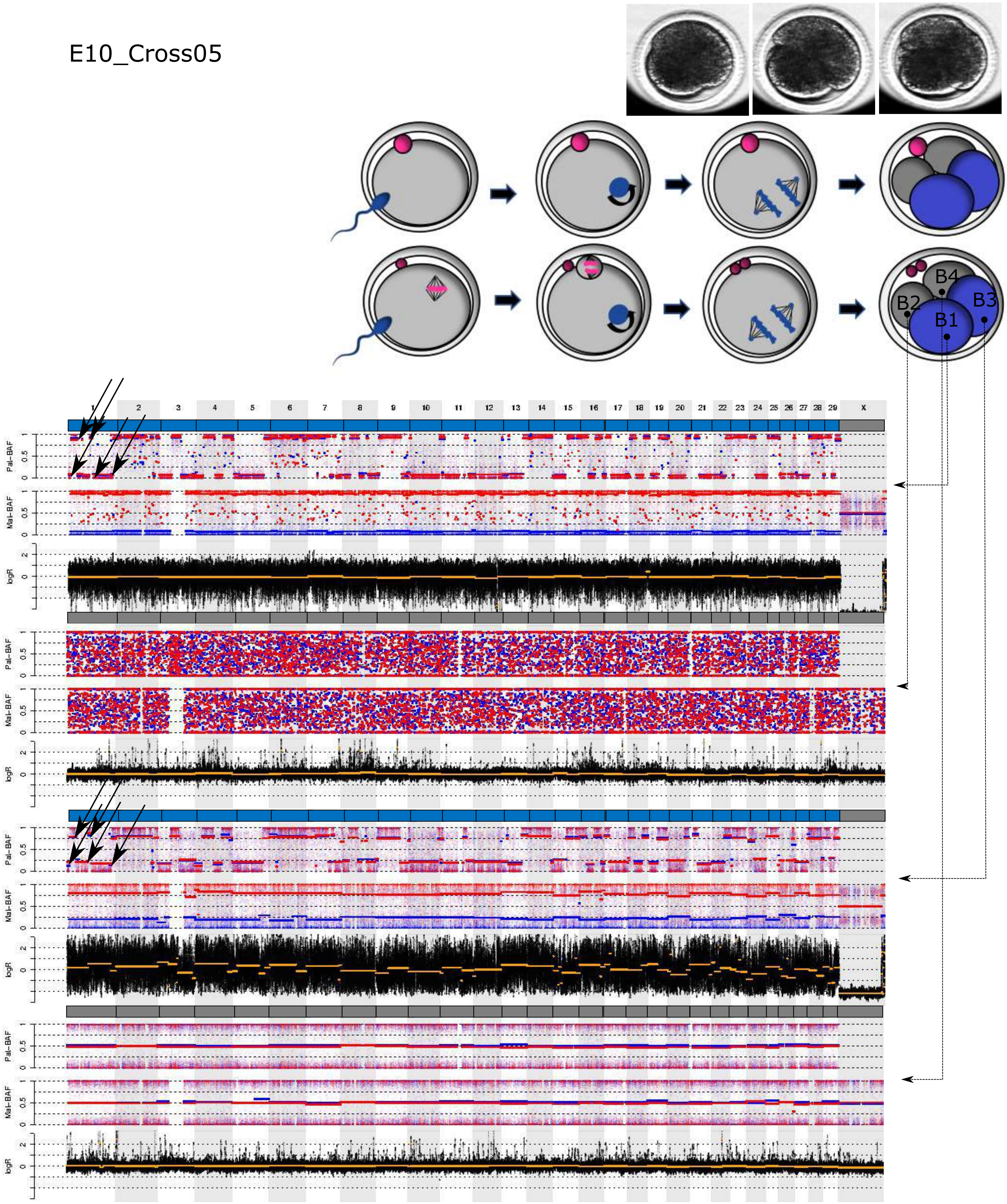

Figure S2B (continued)

### 5. Polyploid embryos

E03_Cross03

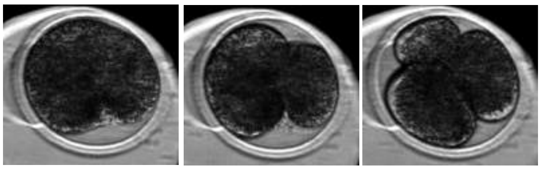

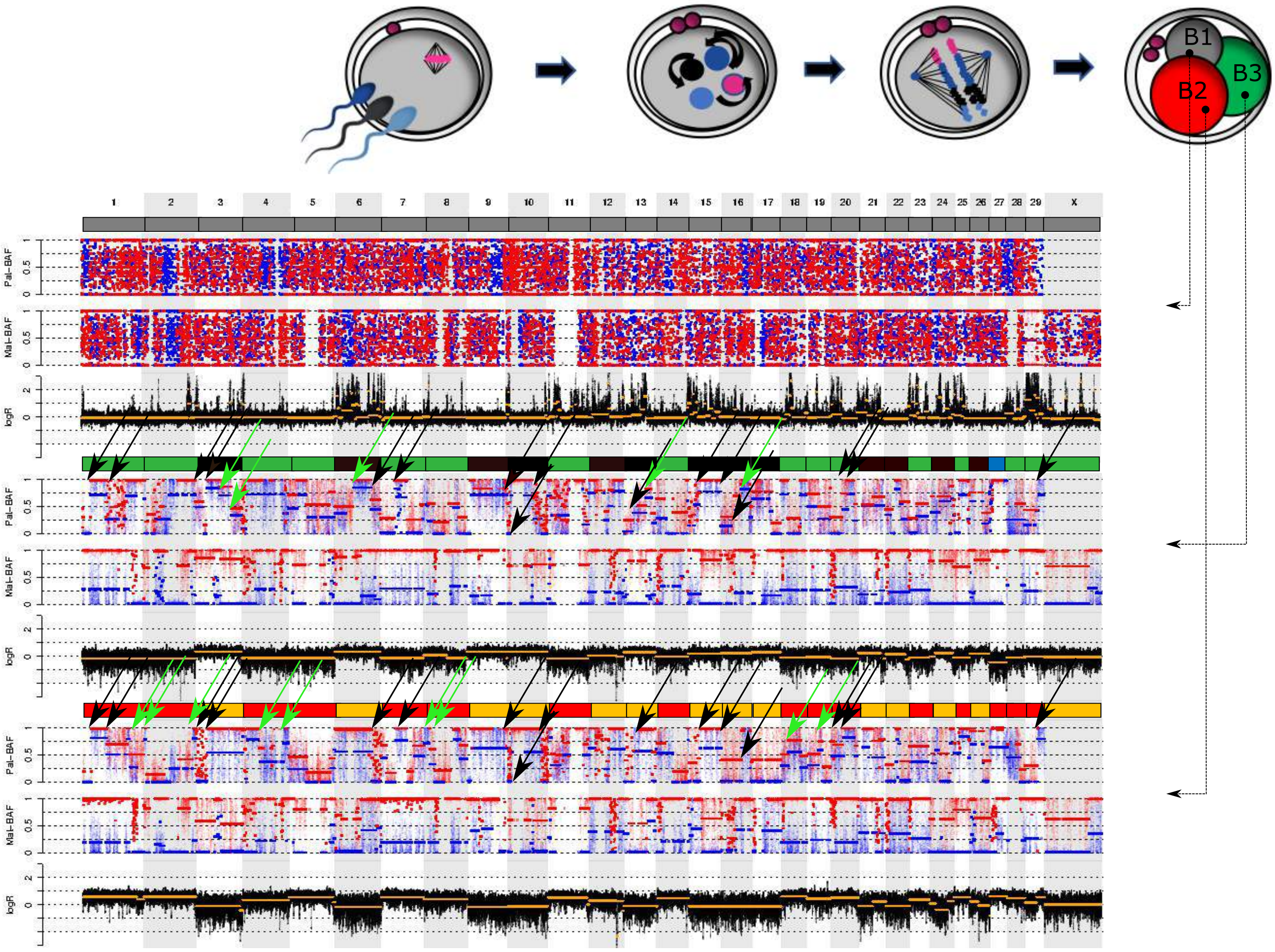

Figure S2B (continued)

E24_Cross12

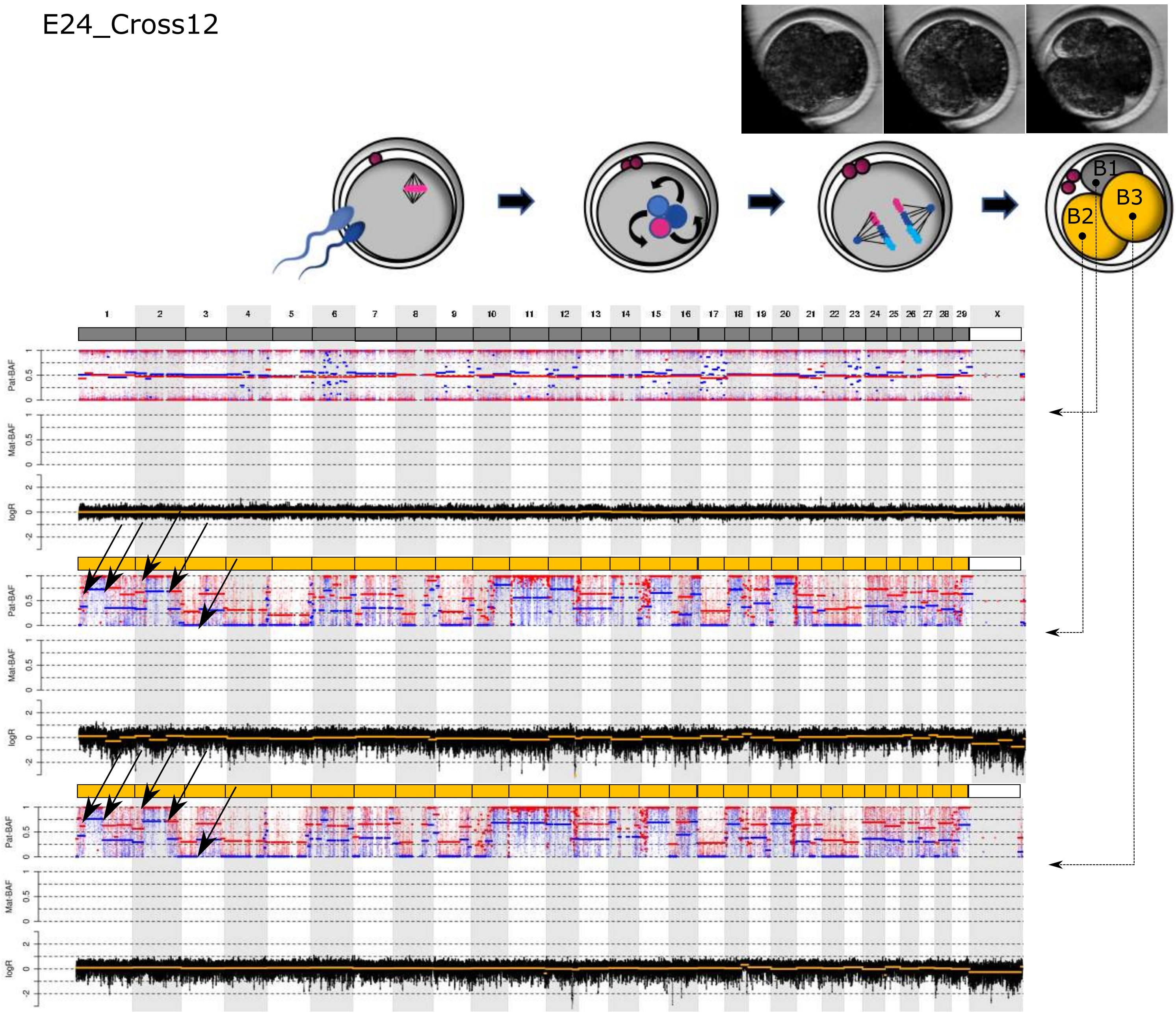

Figure S2B (continued)

E08_Cross04

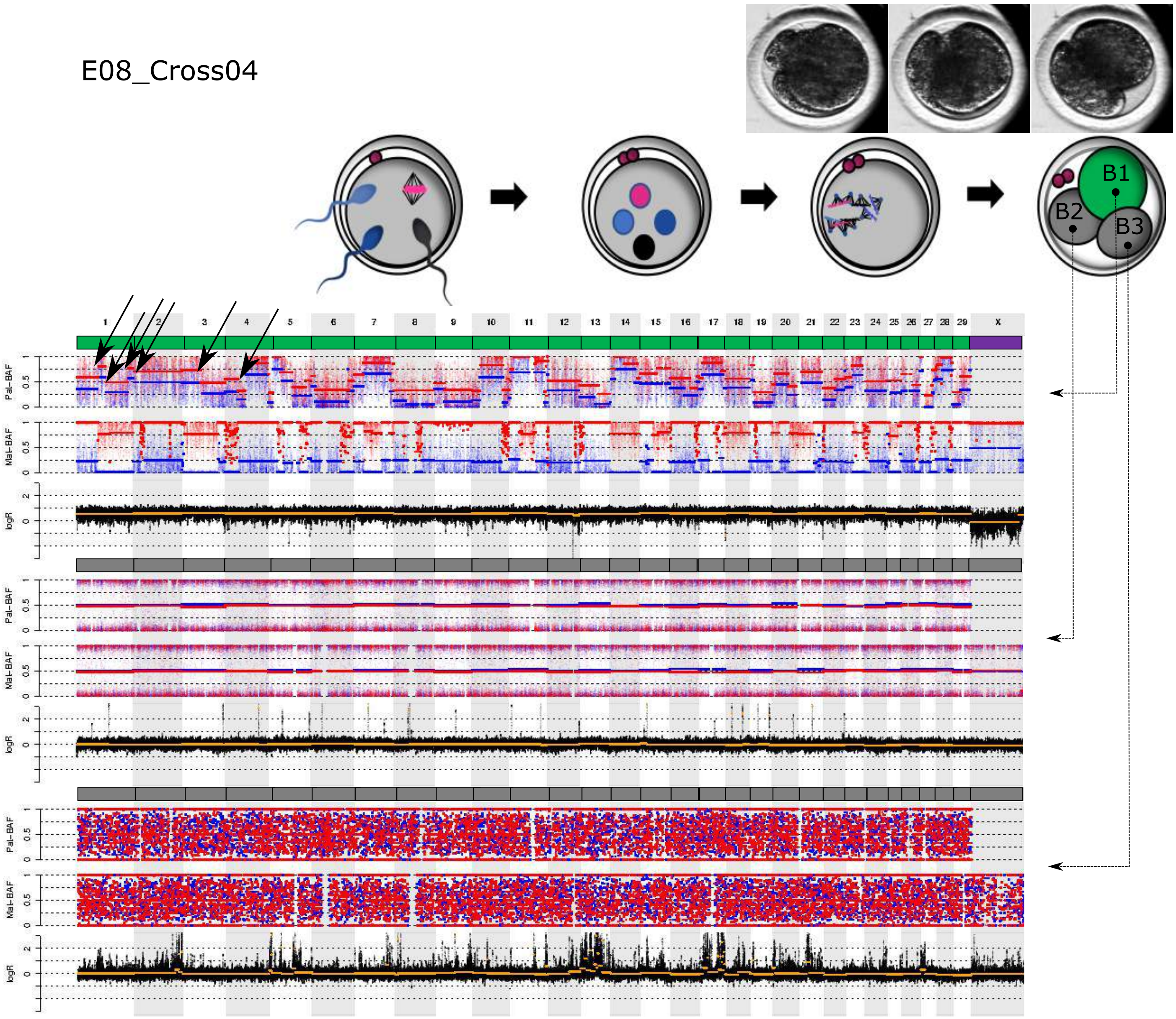

Figure S2B (continued)

### 6. Other profiles

E04_Cross02

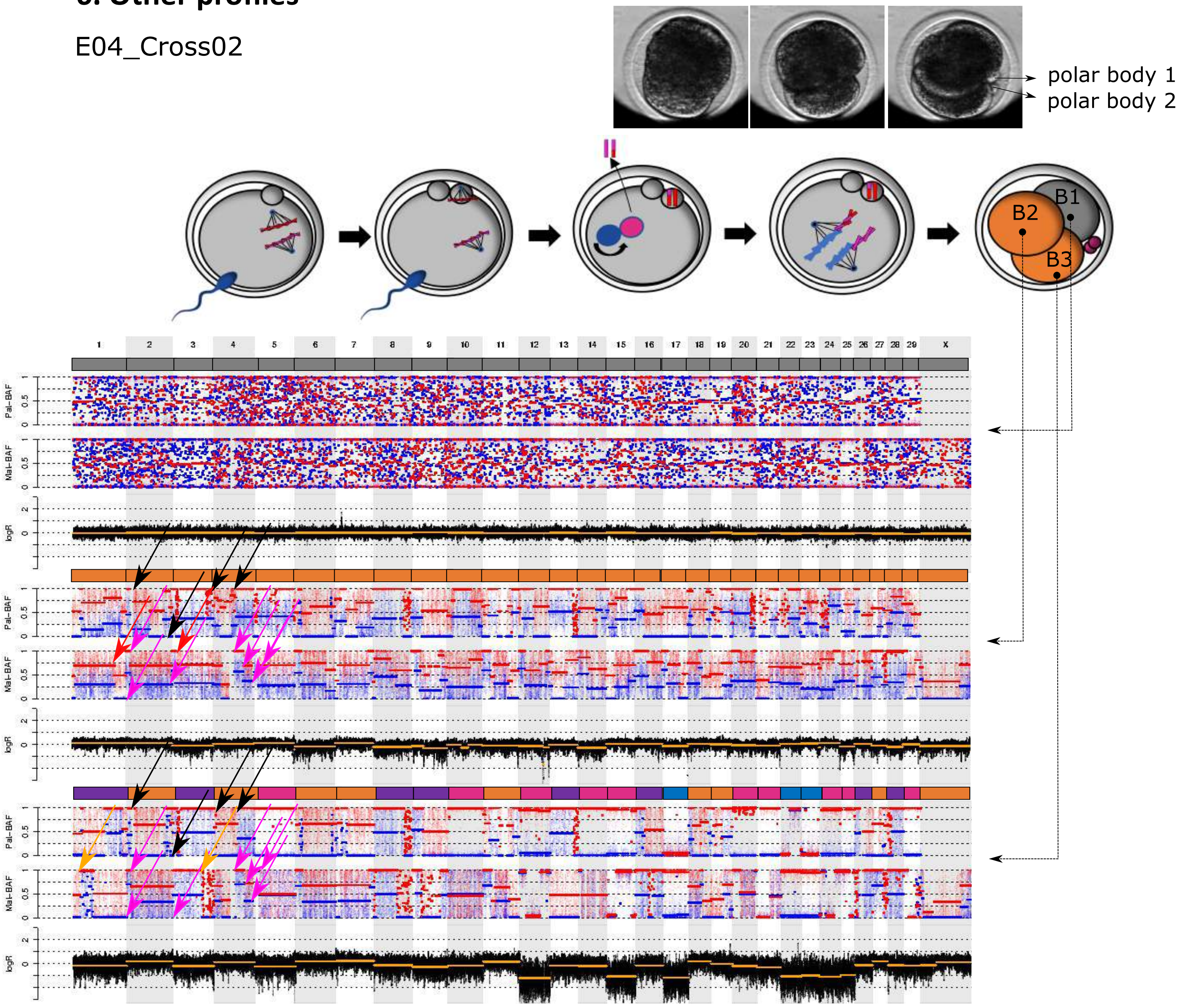

Figure S2B (continued)

E20_Cross08

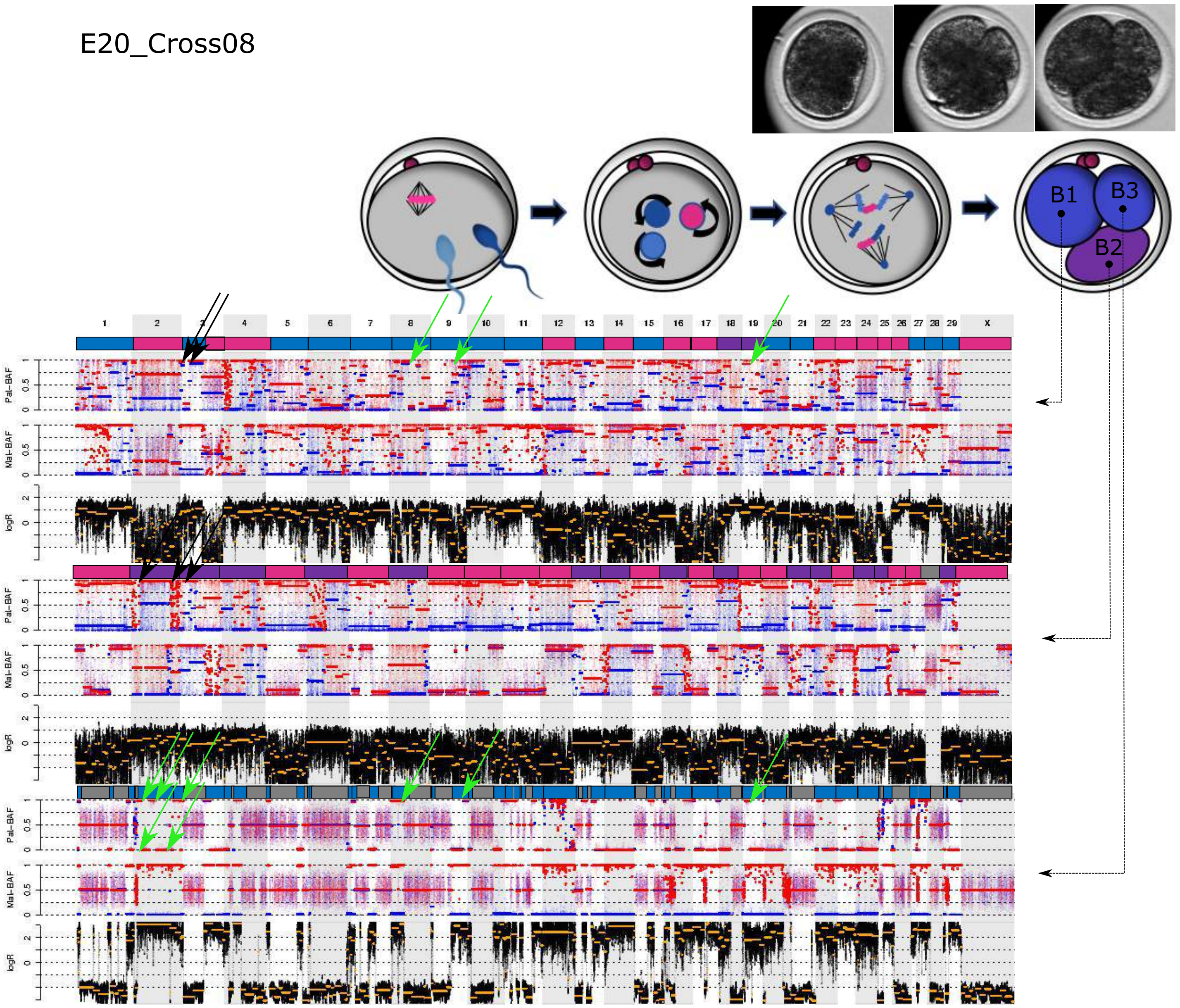

Figure S3

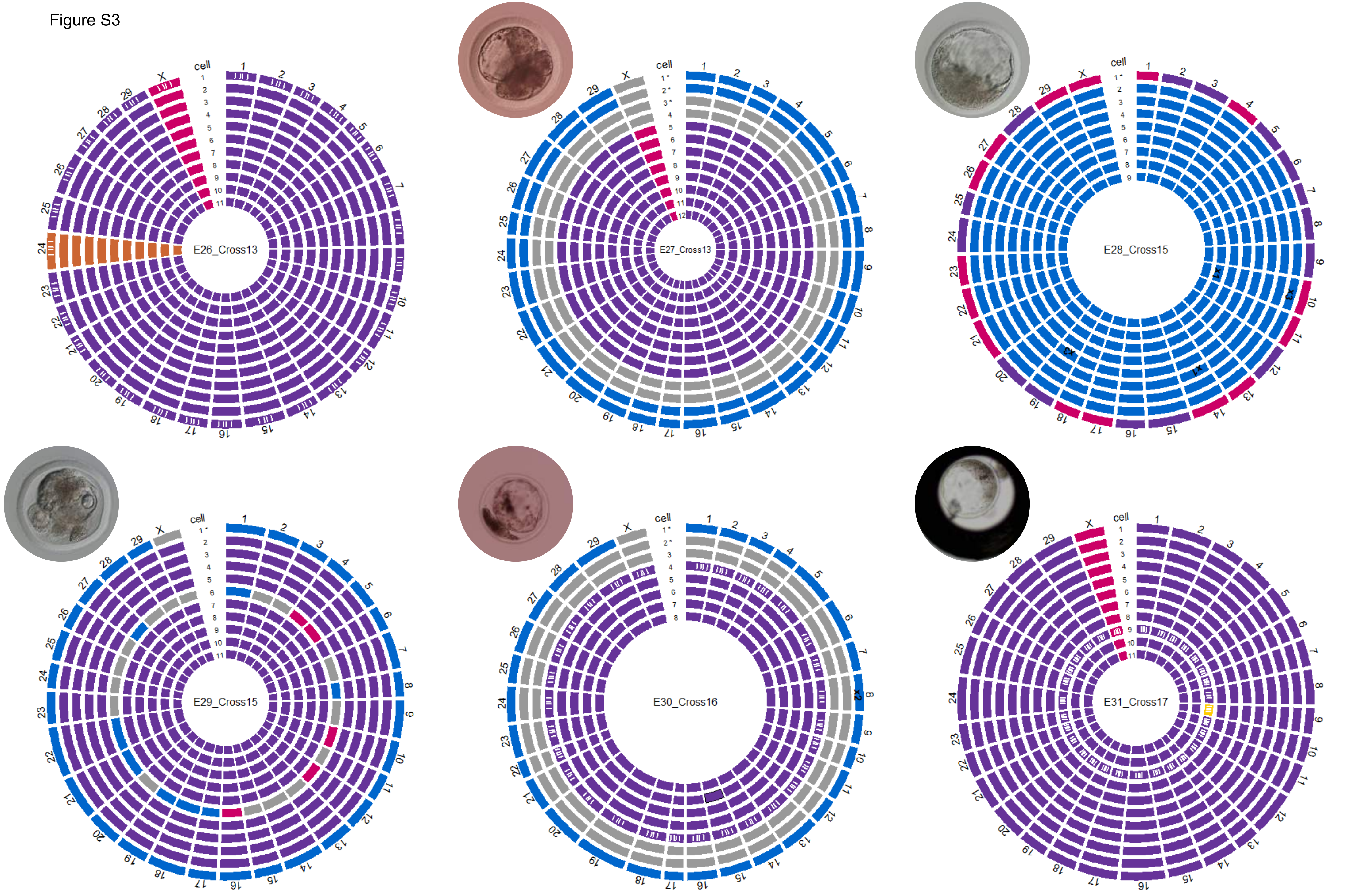


biparental disomy | paternal only | maternal only | digynic trisomy | nullisomy | low quality, presumed biparental disomy | low quality, presumed maternal

low quality, presumed digynic trisomy | low quality, presumed diandric trisomy | * blastomere extruded to periphery | **x1** uniparental chromosomal loss | **x2/x3** uniparental chromosomal gain
